# RBP-Driven RNA Sorting and Local Translation Establish the Molecular Identities of Individual Sensory Axons

**DOI:** 10.1101/2025.02.28.640799

**Authors:** Elizabeth S. Silagi, Ezechukwu Nduka, Maria F. Pazyra-Murphy, Jesus Zuniga Paiz, Shamsuddin A. Bhuiyan, William Renthal, Rosalind A. Segal

## Abstract

Neurons extend long axons that traverse distinct microenvironments, yet how these compartments acquire and maintain specialized molecular identities remains unclear. Here, we use spatial translatomics to define the local axonal translatomes of dorsal root ganglion (DRG) neurons that mediate somatosensation. Translating Ribosome Affinity Purification and RNA sequencing revealed thousands of mRNAs that are preferentially translated within the central axons, peripheral axons, or DRG soma, establishing compartment-specific translational programs. Many of these transcripts encode ion channels, neurotransmitter receptors, and structural proteins that confer distinct electrophysiological, synaptic, and regenerative properties to each axon. Cross-dataset integration with single-cell RNA-seq demonstrated that neuropathic injury elicits highly compartment-specific remodeling of these localized translational programs. We identify polarized RNA regulons coordinated by the RNA-binding proteins (RBPs) SFPQ and SRSF10, which preferentially bind and traffic mRNAs to the peripheral or central axon, respectively. These findings reveal a mechanistic framework in which RBP-guided RNA sorting and local translation establish and dynamically tune subcellular specialization in sensory neurons.

**HIGHLIGHTS:**

- Distinct mRNAs are translated in peripheral or central axons (painseq.shinyapps.io/CompartmentTRAP/).
- Axonal translatomes enable localized regulation of electrophysiology, neuronal plasticity, and regeneration.
- The RBPs, SFPQ and SRSF10, enable mRNA sorting to peripheral and central axons respectively.

## INTRODUCTION

Neurons possess several distinctive and essential features, including the ability to form long-range connections, efficiently propagate electrical signals, and exhibit plasticity in response to multiple stimuli. These capabilities rely on dynamic regulation of gene transcription and translation which together shape the cellular proteome. Transcriptional programs adapt rapidly to changes in the overall neuronal state, including alterations in membrane polarization, metabolic activity, and the inflammatory environment ^1–3^. In contrast to transcription, translation can be regulated at the subcellular level, allowing for localized responses to changes in the microenvironment, such as those occurring at synapses and along the axons and dendrites ^4–10^. As a result, local translation offers an additional layer of spatiotemporal regulation, fine-tuning the neuronal proteome in response to both global and microenvironmental signals.

It was not long ago that scientists believed local translation could not occur in axons. The first evidence of RNA in axons was detected in the 1960s ^11,12^, however, it took another 30 years of technological advances in microscopy for researchers to show that ribosomes were present in axons and capable of translating mRNA ^13,14^. Since 2000, multiple studies have revealed that local protein synthesis is necessary for axon development and regeneration, and proteins transported from the soma are not sufficient to maintain local proteomes ^5,15–17^. Despite this progress, it is not understood how mRNA transport and translation establish local proteomes and govern the specificity of axonal functions. Analysis of local translation in the axons of retinal ganglion cells in the central nervous system (CNS) suggests that translatomes are consistent across distinct axon branches ^18^, which would preclude dynamic regulation of individual axonal proteomes in response to changes in the local microenvironment. However, it is not known whether it is possible for neurons to differentially regulate mRNA transport and translation to establish and modulate distinctive features of individual axons.

In the peripheral nervous system (PNS), sensory neurons of the dorsal root ganglia (DRG), trigeminal ganglia, and nodose ganglia are distinguished by their pseudounipolar morphology, where a single axon stem bifurcates into central and peripheral projections that innervate the spinal cord and peripheral tissues, respectively ^19,20^. In humans, each of these axons can extend up to 1 meter from the cell soma in the ganglion. The peripheral axons of DRG neurons are responsible for dynamically transducing environmental stimuli into electrical signals, while the central axons transmit these signals to the CNS. Both peripheral and central axons generate and propagate action potentials, are myelinated in many DRG subtypes, and exhibit the same microtubule orientation with plus-ends towards the terminals ^20,21^.

Studies of DRG axonal function and composition have historically focused on the peripheral axon as it is more accessible for surgical isolation, injury models, and electrophysiological recordings, while investigations of the central axon are underrepresented ^22^. Therefore, it is not known how the individualized proteomes of central and peripheral axons are decided and established ^23^. The spatially and functionally distinct central and peripheral axon branches make DRG neurons an ideal system for addressing whether subcellular axonal translatomes differ, how any such differences arise, and how they contribute to the compartmentalized demands of encoding, transmitting, and filtering sensory information.

In this study, we identify and compare the spatial translatomes of the two axons of somatosensory neurons. The sodium channel Nav1.8 is expressed in approximately 60% of DRG neurons, including most subtypes involved in pain sensation ^19,24,25^. Using Nav1.8^cre^; L10a-GFP^fl/fl^ mice and a combination of Translating Ribosome Affinity Purification and highly sensitive RNA sequencing (TRAP-Seq) ^26^, we profile the central and peripheral axonal translatomes of somatosensory neurons from microdissected mouse tissues—specifically, 1) lumbar DRG cell bodies, 2) sciatic nerves (peripheral axons), and 3) lumbar dorsal roots (central axons). Our analysis identifies thousands of axon-selective transcripts, including specialized ion channel and receptor isoforms with preferential translation in peripheral and central axons. These findings indicate that translation is an indispensable mechanism for locally modulating axonal physiology within distinct branches of the same neuron. We show that many of the locally translated mRNAs are selectively localized to the central versus peripheral axons, indicating that there is disparate transport and/or stabilization of individual mRNAs in these two axons. Notably, we identify distinct RNA-binding proteins (RBPs) that predominantly bind locally translated mRNAs of the peripheral or central axons, suggesting RBPs enable selective localization of functional RNAs cargoes to individual axons.

Together, our findings offer a comprehensive understanding of how selective RNA transport and translation enable highly specialized subcellular proteomes that establish the spatial organization of neurons in normal physiology and disease states. This resource (http://painseq.shinyapps.io/CompartmentTRAP) provides a wealth of data to investigate how cellular functions are selectively regulated at the local level and highlights how dysregulation of axonal translatomes may contribute to neurodegenerative diseases and disorders.

## RESULTS

### Defining subcellular translatomes in somatosensory dorsal root ganglia neurons

To generate a comprehensive resource of the axonally transported and translated mRNAs in DRG sensory neurons *in vivo*, we performed spatial translatomics using Translating Ribosome Affinity Purification (TRAP) paired with ultra-low methods for RNA-sequencing to account for technical difficulties of extracting high-integrity RNA from axons *in vivo*. The lumbar ganglia (soma), sciatic nerves (peripheral axons), and lumbar dorsal roots and spinal cords (central axons) were microdissected from ∼15 perinatal mice (week 1) containing GFP-tagged ribosomal subunits in Nav1.8+ neuronal cell types (Nav1.8^cre^;L10a-GFP^fl/fl^) (Fig. 1A). Ribosome-associated mRNAs were isolated from tissue lysates using GFP-antibody affinity purification and ultra-low concentration RNA extraction yielding ∼10-100 pg/μL RNA (Fig. S1A,B). We carried out the same procedure on littermate negative control mice (L10a-GFP^fl/fl^) that lacked GFP-tagged ribosomes. Libraries were prepared from low input total RNA samples with SMART-Seq, and subsequently sequenced and aligned to elucidate the distinct central and peripheral axonal translatomes (n=4 sets/condition, each set consisted of 13-17 individual animals) (Tables S1-3) ^27^.

**Figure 1.**
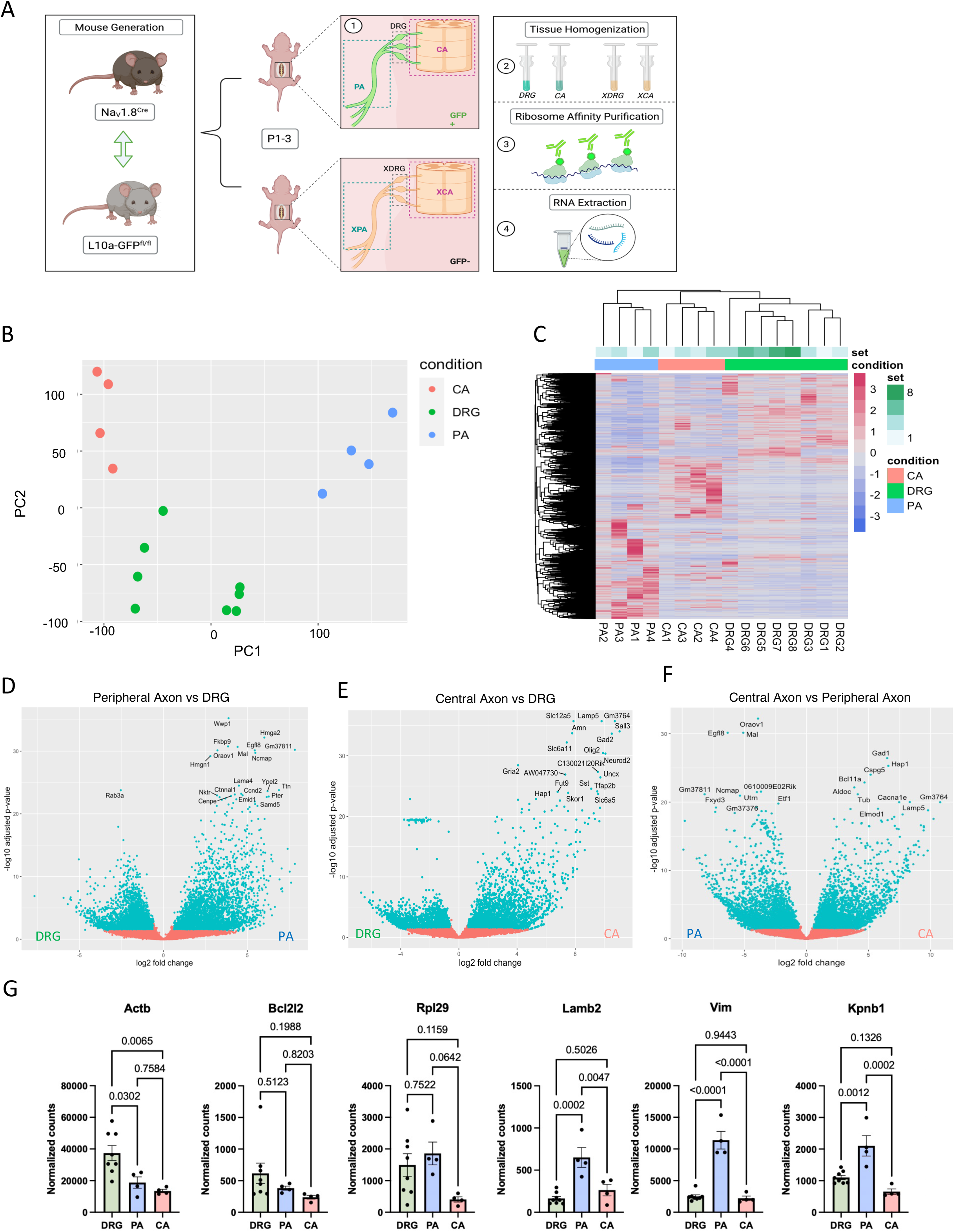
Defining subcellular translatomes in somatosensory dorsal root ganglia neurons. A) Schematic depicting the TRAP-sequencing protocol. Nav1.8Cre/+ mice were crossed to L10a-EGFPfl/fl mice to generate Nav1.8Cre;L10a-EGFPfl/fl mice and L10a-EGFPfl/fl negative control mice. In these mice, Nav1.8-Cre drives expression of an EGFP-tagged ribosomal subunit, L10a, selectively in a set of DRG neurons. Tissue microdissections were performed in mice aged P1-3 to collect lumbar dorsal root ganglia (soma compartment), lumbar dorsal roots/ spinal cord (central axon compartment), and sciatic nerves (peripheral axon compartment). Using the Translating Ribosomal Affinity Purification (TRAP) protocol, ribosomal-associated mRNAs were isolated from tissue lysates using an anti-EGFP antibody immunoprecipitation followed by RNA extraction using filter columns. B) Principal component analysis of DRG soma, peripheral axon, and central axon samples (n=4 peripheral and central axon samples, n=8 DRG soma samples). C) Heatmap showing normalized gene expression within the DRG soma, peripheral axon, and central axon TRAP datasets for all genes with expression ≥ 5 counts. D-F) Volcano plots demonstrating differential gene expression between the TRAP datasets comparing peripheral axons vs. DRG soma (D), central axons vs. DRG soma (E), and central vs. peripheral axons (F) (p.adj ≤ 0.05; fold-change ≥ 1.5). G) Bar graphs depicting the normalized gene expression of select mRNA transcripts (with known expression in axons) within the DRG soma, peripheral axon, and central axon compartments. Statistical significance determined by two-way ANOVA with Tukey’s multiple comparisons test (p-value ≤ 0.05). Data are represented as mean ± SEM.

Principal component analysis and pairwise comparisons demonstrate that the DRG soma samples cluster together across the multiple samples, while central and peripheral axon samples cluster distinctly from the soma and from each other (Fig. 1B). This distinct clustering indicates that the transcripts were highly consistent and specific for each compartment. Heatmap visualization with hierarchical clustering of ∼18,000 normalized genes revealed a clear segregation of samples by condition and distinct translational profiles across DRG soma, peripheral axon, and central axon samples (Fig. 1C). Volcano plots demonstrating the significance and fold-change between compartments for all genes indicated that both peripheral and central axons expressed many genes that were enriched (padj. ≤ 0.05; F.C. ≥ 1.5) over the DRG compartment, demonstrating substantial axonal translation in DRG sensory neurons in vivo (Fig. 1D,E). Moreover, the volcano plot comparing central and peripheral axon compartments indicates that large numbers of distinct transcripts are translated in the peripheral versus central axons, demonstrating the subcellular specificity with which DRG neurons control local translation to regulate spatially distinct neuronal functions in DRG soma and peripheral and central axons (Fig. 1F).

Interestingly, previously characterized axonally translated genes such as *Actb* (β-actin), *Bcl2l2* (Bclw), *Rpl29* (60s ribosomal protein L29), *Lamb2* (Laminin β2), *Vim* (Vimentin), and *Kpnb1* (Importin β1) are all enriched in the peripheral axon compartment and have lower expression in the central axon (Fig. 1G). This suggests that the characterization of axonal translatomes in scientific literature mirrors DRG peripheral axons, while the translational landscape of DRG central axons may differ from the prior data. Thus, defining what is locally translated in both peripheral and central DRG axons is critical for understanding how compartment-specific protein synthesis contributes to somatosensory information transfer.

### Central and peripheral DRG axons have distinct translatomes

Our analysis reveals that the peripheral and central axonal translatomes are remarkably different from one another (Fig. 2A;Table S4). In total, the DRG axonal translatome contains roughly 13,000 genes, of which 5,960 genes are shared between the two axons, 3,379 are enriched in central axons and 3,420 are enriched in peripheral axons (Fig. 2B). Gene ontological (GO) functional enrichment analysis of the genes shared between the axonal compartments reveals local translation of transcripts related to autophagy, RBP complex formation, ubiquitin-dependent proteasomal mechanisms, mitochondrial function, and ribosome biogenesis in both axons (Fig. 2C; Table S5). Many of these transcripts are also translated in the cell soma; ∼60% of the transcript translated in peripheral axons were also translated in soma, and 72% of the transcripts identified in the TRAP-Seq of central axons were translated in the soma as well (Fig. S2A-C), and these ubiquitous components are also related to ubiquitin-dependent proteasomal degradation, autophagy, ribonucleoprotein complexes, and organelle function (e.g. mitochondria, Golgi, endoplasmic reticulum, and ribosomes) (Fig. S2D,E).

**Figure 2.**
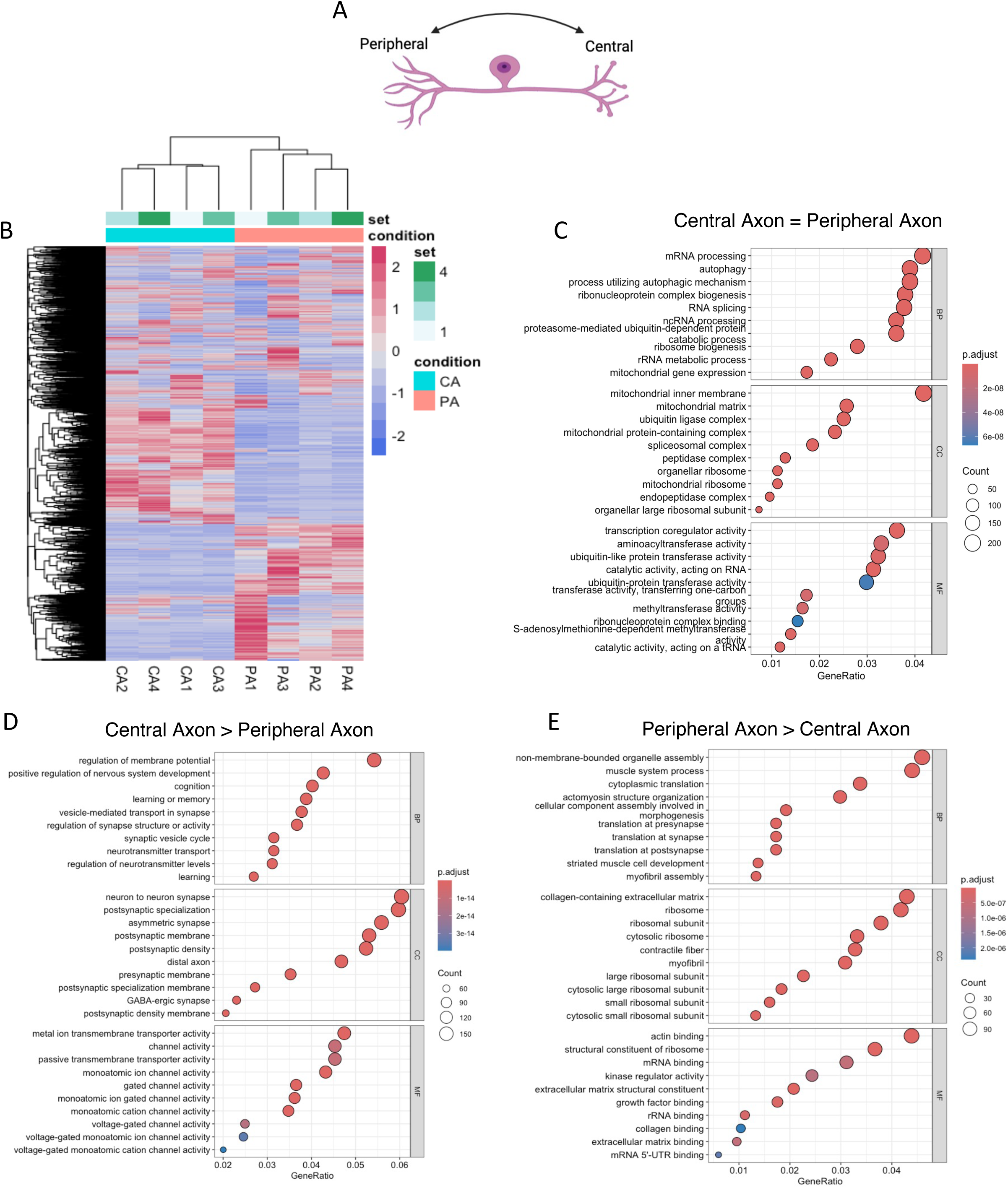
Central and peripheral DRG axons have distinct translatomes. A) Schematic depicting the translatomic gene sets compared in Figure 2 (i.e. central axon vs. peripheral axon). B) Heatmap highlighting clusters of genes with 1) similar expression across central and peripheral axon compartments, 2) higher expression in the central axon compartment, and 3) higher expression in the peripheral axon compartment (n=4 samples/group). B) Gene ontology (GO) enrichment analysis visualized as a dot plot, where each dot represents a significantly enriched biological process (BP), cellular component (CC), and molecular function (MF) from the translatomic gene set shared between both axons (p.adj ≥ 0.05; abs[F.C] ≤ 1.5). C) Dot plot showing GO terms enriched in the central axon geneset (p.adj ≤ 0.05; F.C ≥ 1.5) D) Dot plot showing GO terms enriched in the peripheral axon dataset. (p.adj ≤ 0.05; F.C ≤ −1.5).

Functional annotation analysis of the axon-specific transcripts that are not shared with the DRG soma, identified local translation of components related to synaptic specialization in the central axons, while peripheral axon-specific transcripts encoded proteins related to cytoskeletal structure and contractility (Fig. S2F,G; Tables S6,7). The 3,379 genes comprising the central axon translatome were enriched for transcripts required for neuronal excitability, synaptic organization, and neurotransmission (Fig. 2D; Table S8). Interestingly, the top enriched terms identified prominent translational programs dedicated to maintaining both presynaptic and postsynaptic function. Genes implicated in presynaptic assembly included *Nrxn1, Cbln1/2, Pclo, and Efnb3*, indicating that there is local synthesis of adhesion molecules and active-zone scaffolds that organize vesicle docking and release sites for neurotransmission.

Furthermore, axonal translation of voltage-gated calcium channel components and SNARE proteins (*Cacng2/4/8, Stx1b, Vamp2*) highlight local modulation of presynaptic transmission efficacy. In addition to these presynaptic components, local synthesis of postsynaptic scaffolding and receptor-associated proteins such as *Grip1/2, Dlg2/3, Gphn, Nptxr*, and *Arc* suggests that DRG central axons also have the capacity to remodel postsynaptic machinery, and thereby alter synaptic activity in response to spinal cord interneurons or descending circuits. Based on the classes of locally translated receptors, we can surmise that local modulation of presynaptic DRG axon terminals occurs through glutamatergic (*Grin/Gria/Grm/Grik*), GABA-ergic (*Gabra/Gabrb*), endogenous opioid (*Oprk/Oprl*) and serotonergic circuits (*Htr*), which aligns with the current known mechanisms of pain modulation in the dorsal horn. Collectively, these findings suggest that DRG central axons are not passive conduits of sensory information but are instead translationally competent compartments that bidirectionally regulate pain signaling through ongoing synthesis of receptors, ion channels, and signaling proteins.

In contrast, GO analysis of 3,420 transcripts locally translated in the peripheral axons of DRG neurons revealed the transcripts were enriched for processes related to translational capacity, cytoskeletal organization, and myelination-associated components (Fig. 2E; Table S9). The enriched mRNAs were strongly associated with ribosomes(*Rpl/Rps*) and cytoplasmic translation (*Eif/Etf*), indicating abundant local synthesis of ribosomal proteins and translation factors that may sustain high levels of protein turnover at distal sites in the axon. Furthermore, terms such as actomyosin structure organization and myofibril assembly pointed to active regulation of the axonal cytoskeleton and contractile elements that support axonal integrity, transport, and regeneration. For example, local translation of the Rho GTPase signaling pathway controlling myosin contractility (*Rhoa, Rock1/2, Cdc42bpa*), actin cytoskeletal scaffolding proteins (Iqgap1-3), and non-muscle myosins (Myh9/Myh14) collectively support local control of growth-cone traction and axon shaft stability. Lastly, local translational control of components required for axon–Schwann cell and axon-ECM (Pak2,*Itgb1, Sdc4, Phldb2*) interactions help explain how peripheral axons sense and adhere to ECM and Schwann-cell surfaces and dynamically remodel contacts during growth and repair. Together, these findings highlight the importance of autonomous ribosomal repair, structural remodeling, and myelin maintenance within the long axons of the peripheral nervous system. This suggests that peripheral axons maintain a translationally rich environment specialized for the regenerative capacity of such axons.

### Cross-dataset integration reveals compartment-specific translational remodeling during neuropathy

Peripheral nerve injury induces a robust reprogramming of the axonal transcriptome and proteome, with increased axonal protein synthesis and dynamic changes in membrane protein composition, and often leads to peripheral neuropathy^28^. However, the mechanisms governing these adaptations are not well understood. In chemotherapy-induced peripheral neuropathy (CIPN), impaired axonal transport of SFPQ-containing RNA granules has been linked to reduced expression of key neuronal survival factors, implicating a connection between axonal mRNA regulation and axon degeneration ^29,30^. To determine whether locally translated mRNAs in distinct neuronal domains are selectively altered during neuropathic injury, we performed a cross-dataset comparison between our compartment-resolved TRAP translatomes and a published single-cell RNA-seq dataset of DRG neurons treated with paclitaxel (Fig. 3A; Table S10). In these previous studies, mice were injected intraperitoneally with 4mg/kg paclitaxel every other day for six days (four injections total), and transcripts altered in each of the DRG cell subtypes following paclitaxel treatment were identified ^28^. Across all DRG neuron subtypes, ∼1–7% of transcripts present in our datasets were significantly changed by paclitaxel (Fig. 3B). The proportion of affected transcripts was highest for those translated in the soma and those shared among multiple compartments. Individual analysis of paclitaxel-sensitive genes in Nav1.8+ DRG subtypes, including Mrgprd+ non-peptidergic nociceptors (NPs), Tac1+/Gpx3+ peptidergic nociceptors (PEP1), Tac1+/Hpca+ peptidergic nociceptors (PEP2), Nefh+ Aβ low-threshold mechanoreceptors (LTMRs) (NF1), and Sst+ pruriceptors (SST) further demonstrate that paclitaxel treatment causes changes across all domains of pain-sensing neurons (Fig. 3C). However, these analyses also suggest that paclitaxel has a greater impact on transcripts in peripheral axon translatome than the central axon.

**Figure 3.**
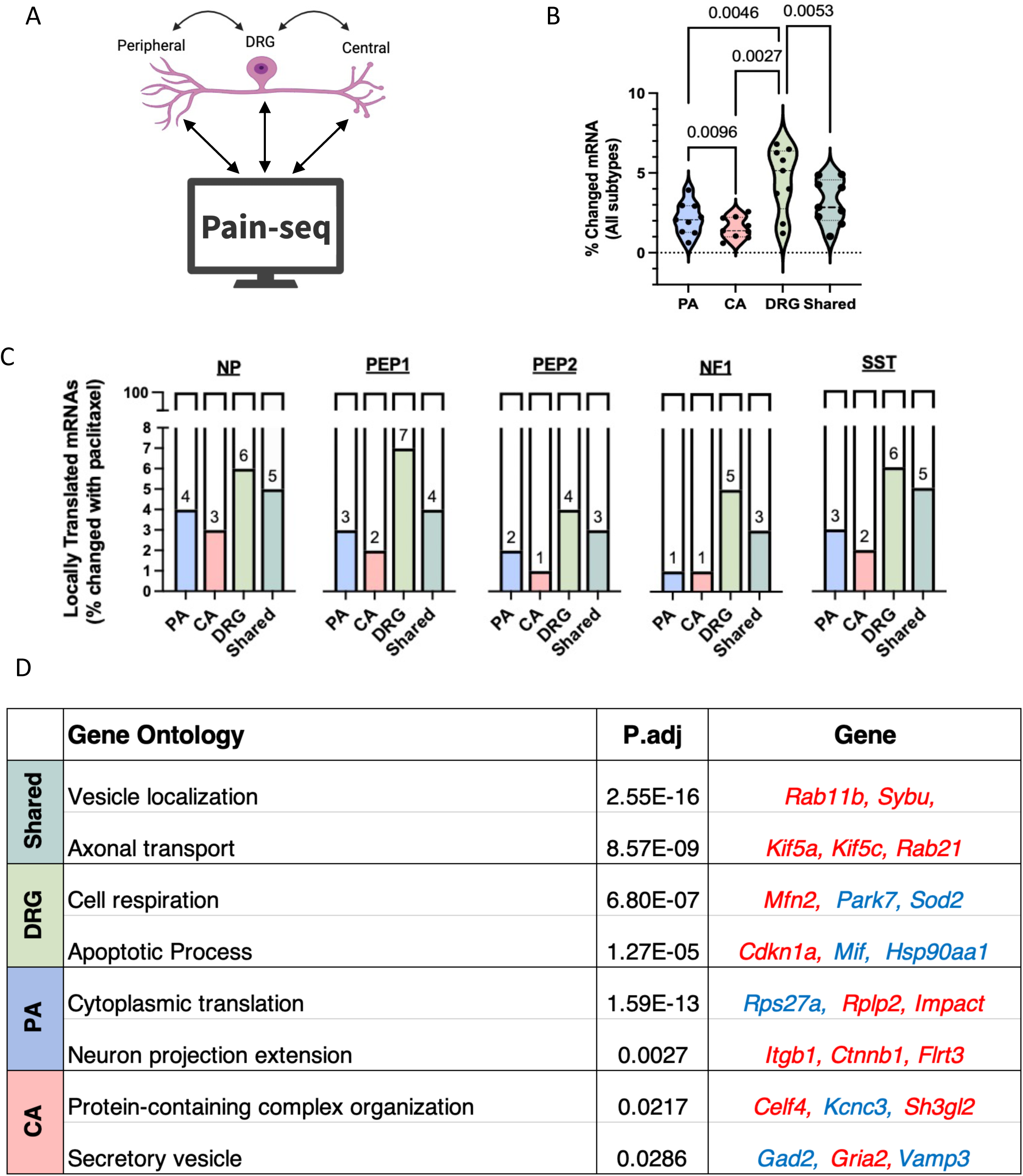
Cross-dataset integration reveals compartment-specific translational remodeling during neuropathy. A) Schematic depicting the cross-dataset integration performed in Figure 3 (i.e. Paclitaxel-treated DRG scRNA-seq (Pain-seq) vs. TRAP datasets from all tissues). B) Bioinformatic analysis showing the percentage of locally translated mRNAs from the TRAP-seq datasets (peripheral axon, central axon, DRG soma, and shared soma + axon) that are differentially expressed in the Pain-seq datasets (following treatment with paclitaxel). Each datapoint represents a different DRG subtype identified via scRNA-seq. B) Percentage of locally translated mRNAs in the TRAP-seq peripheral axon, central axon, soma, and shared datasets that are differentially expressed in Pain-seq datasets following treatment with paclitaxel, specifically in Nav1.8+ DRG subtypes (NP, PEP1, PEP2, NF1, and SST). C) GO enrichment analysis showing significantly enriched GO terms and select locally translated genes from each tissue compartment that are differentially expressed following paclitaxel treatment (p.adj ≤ 0.05). Red: mRNAs that are upregulated following paclitaxel treatment. Blue: mRNAs that are downregulated following paclitaxel treatment.

To identify the biological programs most strongly impacted within each subcellular compartment, we performed GO enrichment analysis on mRNAs whose overall expression levels change with paclitaxel and are locally translated in one or another domain (Fig. 3D). Paclitaxel affects multiple mRNAs that are “shared” in soma and axonal translatomes; these locally translated mRNAs are involved in axonal transport and vesicle localization, such as *Rab11b, Sybu*, and *Kif5* family motors, implying that paclitaxel reconfigures intracellular trafficking pathways that operate across neuronal compartments. Furthermore, the somatic compartment upregulated pathways related to cell respiration and apoptotic signaling, including *Park7, Sod2*, and *Cdkn1a*. Paclitaxel alters many mRNAs that are predominantly translated in the peripheral axon compartment, including components involved in cytoplasmic translation, environmental sensing, and neuronal projection extension, such as *Itgb1, Ctnnb1*, and *Flrt3*, indicating enhanced remodeling of local growth and structural pathways. In contrast, paclitaxel-sensitive mRNAs that are translated in the central axon include transcripts linked to protein-complex assembly and synaptic/ secretory function, such as *Celf4, Kcnc3, Sh3gl2*, and *Gad2*, reflecting changes in neurotransmitter release and presynaptic physiology. Together, these analyses reveal that paclitaxel-induced neuropathy evokes extensive remodeling in DRG neurons, with each neuronal domain enacting a distinct adaptive or maladaptive response during the development of neuropathic pain.

### DRG axons locally translate specialized ion channels and membrane-bound receptors

While analysis of the initial axonal translatomes demonstrates that the translatomes of the two axons differ greatly from one another, some RNAs identified in the bulk sciatic nerve, dorsal root, and spinal cord tissues may not be derived from Nav1.8+ sensory axons, but rather from local glia, epineural cells, and neuronal cell bodies in the spinal cord. For example, compared to the DRG compartment, the central translatome exhibited high levels of *Olig2*, a marker of oligodendrocytes, and the peripheral translatome had high levels of *Ncmap*, a marker of myelinating Schwann cells. To distinguish which of the thousands of transcripts were definitively derived from Nav1.8+ sensory neurons, rather than other cells, we determined the transcripts with significantly higher expression in TRAP-Seq of sciatic nerves, DRG or dorsal roots in Nav1.8-GFP mice as compared to negative controls of TRAP-seq from the same locations in mice lacking L10a-GFP (p.adj ≤ 0.05) (Fig. S3A-E) ^31^. After bioinformatically filtering out the background transcripts with high expression in negative controls, the specific translatomic gene lists were refined to include 549 transcripts from the soma, 304 from the peripheral axons, and 225 from the central axons (Table S11). This method of stringent background filtering established higher confidence cohorts of locally translated mRNAs within DRG soma and peripheral and central axons.

Our background filtered list of locally translated mRNAs confirmed that central and peripheral axons locally translate markedly distinct proteins, including many specialized synaptic receptors and ion channels required for sensory processing. For example, local translation of serotonin receptors, *Htr3a* and *Htr5a,* in the central axons suggest serotonin binding may be an important component of descending modulation of nociceptive inputs within the primary somatosensory neurons (Fig. S3F). Several genes encoding ion channel proteins were identified in this central axon gene set, including *Scn11a,* which encodes Nav1.9. The sodium channel Nav1.9 is selectively expressed in DRG neurons and point mutations in this channel protein can cause either hyperexcitability and chronic pain or chronic insensitivity to pain ^32^. Several distinct ion channels important for action potential propagation were identified in the higher confidence peripheral axon dataset, such as *Hcn1* and *Scn2a* (Nav1.2) (Fig. S3G). We also identified transcripts encoding the calcium channel, Catsper*2*, which was initially identified as a voltage-dependent, Ca^2+^ selective, pH-sensitive ion channel highly expressed in sperm, as well as two ancillary components of this channel (Catsperd and Catsperζ) ^33^. Thus, several components that are locally translated in the peripheral axons also play critical roles in membrane physiology and nociception and indicate that regulated axonal translation in both central and peripheral axons may modify pain responses.

To verify our findings of local and selective translation in the central or peripheral axons, we focused on several channels and receptors identified in our high confidence, background filtered screen (Table S12). Puromycin was intraperitoneally injected into mice for 20 minutes to label nascent protein chains in the dorsal root ganglia, sciatic nerves, and dorsal roots (Fig. 4A). Proximity labeling (Puro-PLA) enabled the visualization of specific nascent proteins via proximity of antibodies bound to puromycin and the translating protein of interest (Fig. S4). To make sure that we are analyzing nascent peptides within the axons analyzed in our TRAP experiments, we focused on PLA puncta that are within the Nav1.8 labeled axons. PLA visualization and quantification verify that individual mRNAs identified by TRAP-Seq as being preferentially translated in the peripheral axons of the sciatic nerve (*Hcn1* and *Scn2a*) are indeed translated there, and that we find very little translation in the central axons of the dorsal roots (Fig. 4B-C’’). In contrast, our analysis indicates that there is extensive translation of *Htr3a* and *Scn11a* in central axons, with little or no translation taking place in the peripheral axons (Fig. 4D-E’’).

**Figure 4.**
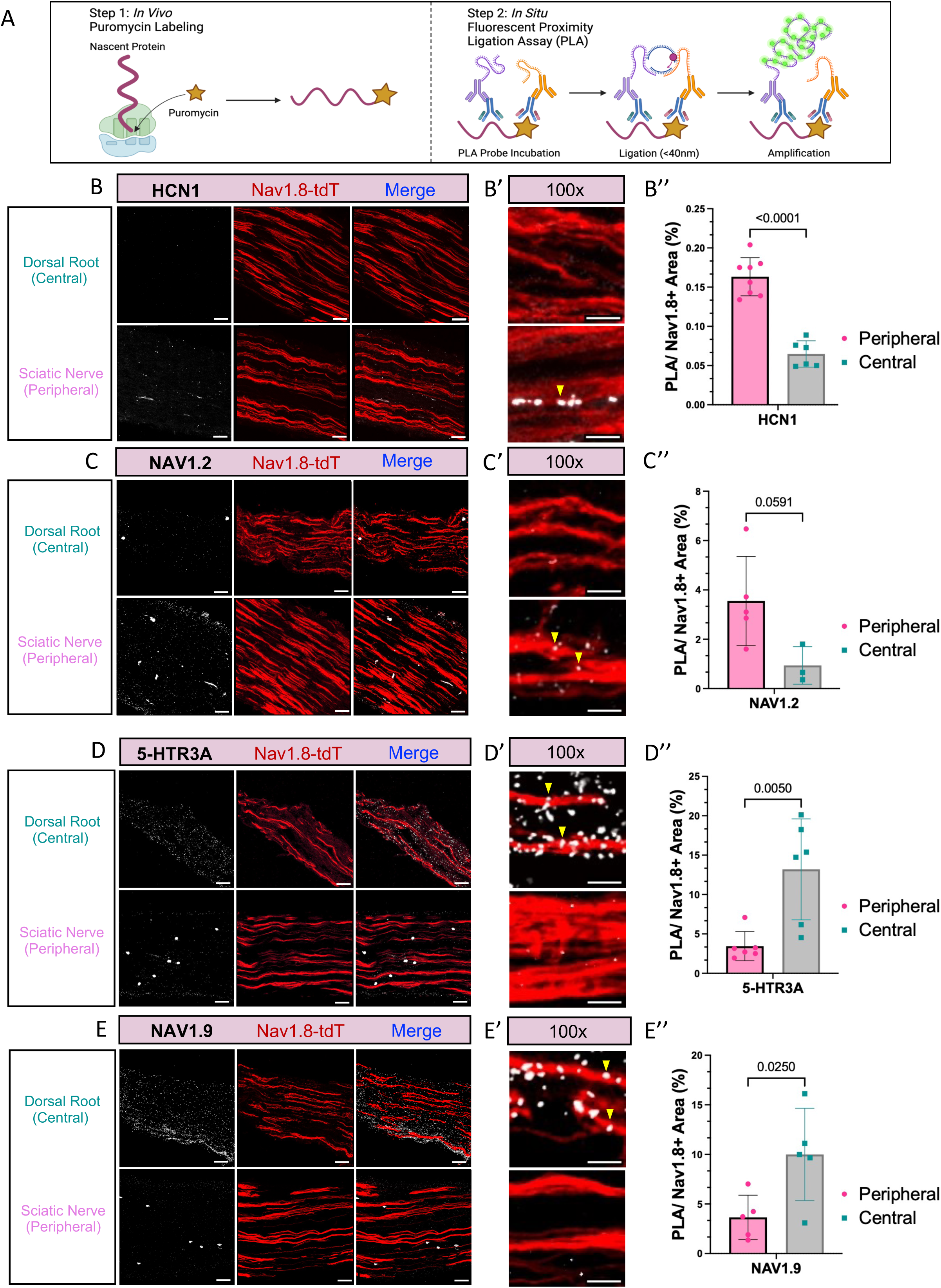
Diverse membrane-associated ion channels are locally translated. A) Puromycin proximity ligation assay (Puro-PLA) enables *in situ* visualization of nascent protein in mouse tissues. Step 1: Puromycin injected into mice incorporates into the C-terminus of translating proteins, labeling nascent peptide chains. Step 2: PLA probes bind two targets of interest (i.e. Puromycin and ion channel) and enable amplification of a fluorescent signal if the targets are within 40 nm. B-E) Representative 20x fluorescent images of HCN1 (B), NAV1.2 (C), 5-HTR3A (D), and NAV1.9 (E) PLA puncta and TdTomato-labeled Nav1.8+ (Nav1.8-tdT) axons in dorsal roots and sciatic nerves, representing DRG central and peripheral axons, respectively. Scale bar = 40 μm. B’-E’) Higher magnification images demonstrating the location and abundance of HCN1 (B’), NAV1.2 (C’), 5-HTR3A (D’), and NAV1.9 (E’) puncta within Nav1.8-tdT+ axons in the dorsal root and sciatic nerve. Scale bar = 10 μm. B’’-E’’) Quantification of the percent area occupied by HCN1 (B’’), NAV1.2 (C’’), 5-HTR3A (D’’), and NAV1.9 (E’’) PLA puncta within Nav1.8-tdT+ axons in dorsal roots and sciatic nerves. Statistical significance determined by two-way ANOVA with Tukey’s multiple comparisons test and multiple unpaired t-tests (p-value ≤ 0.05). Data are represented as mean ± SEM. n=1-2 sciatic nerves samples/mouse and 1-3 dorsal roots samples/mouse from 4-5 mice.

Together, the results seen with nascent-chain puromycin labeling (Puro-PLA) validate the specificity and sensitivity of our TRAP-seq approach and demonstrate that the background-filtered translatomes constitute a powerful resource for discerning axonally synthesized proteins. Moreover, our data show that discrete cohorts of ion channels and receptors are synthesized locally within each axonal compartment, supporting the conclusion that peripheral and central branches possess distinct translational identities that can contribute to their specialized physiological roles in nociceptive signaling.

### mRNAs encoding locally translated proteins are selectively targeted to axons

For the identified proteins to be translated in either axon, the corresponding mRNA must both be transported to that axon and engage with translational machinery at that location. To determine whether the distinct translatomes of the central and peripheral axon reflect differential localization of the mRNA transcripts, or whether these mRNAs are similarly localized to both axons but are selectively translated at one site, we utilized *in vivo* RNAscope fluorescent *in situ* hybridization (FISH) to evaluate the subcellular localization of select mRNAs. Single molecule RNAscope demonstrated the mRNA expression pattern for mRNAs encoding *Hcn1, Scn2a, Htr3a, Scn11a, Catsper2*, and *Trpv1* within the DRG soma, sciatic nerves (peripheral), and dorsal roots (central) (Fig. 5A-D’, Fig. S5). Again, we focused on RNAscope puncta that are within the Nav1.8 labeled axons, or within the Nav1.8 cell bodies in the DRGs. Using particle analysis, we found that *Hcn1, Scn2a,* and *Catsper2* transcripts have significantly more puncta per area in Nav1.8+ peripheral axons compared to central axons, indicating that these transcripts are both selectively localized and selectively translated in peripheral axons (Fig. 5E,F). In contrast, *Htr3a* and *Scn11a* have significantly more puncta per area in Nav1.8+ central axons than in peripheral axons (Fig. 5G,H). As expected, all these mRNAs were also localized to the Nav1.8+ cell bodies of DRG neurons. These data indicate that the distinct axonal translatomes in the central and peripheral axons predominantly reflect differential transport and localization of the mRNAs from the cell body to one axon or the other.

**Figure 5.**
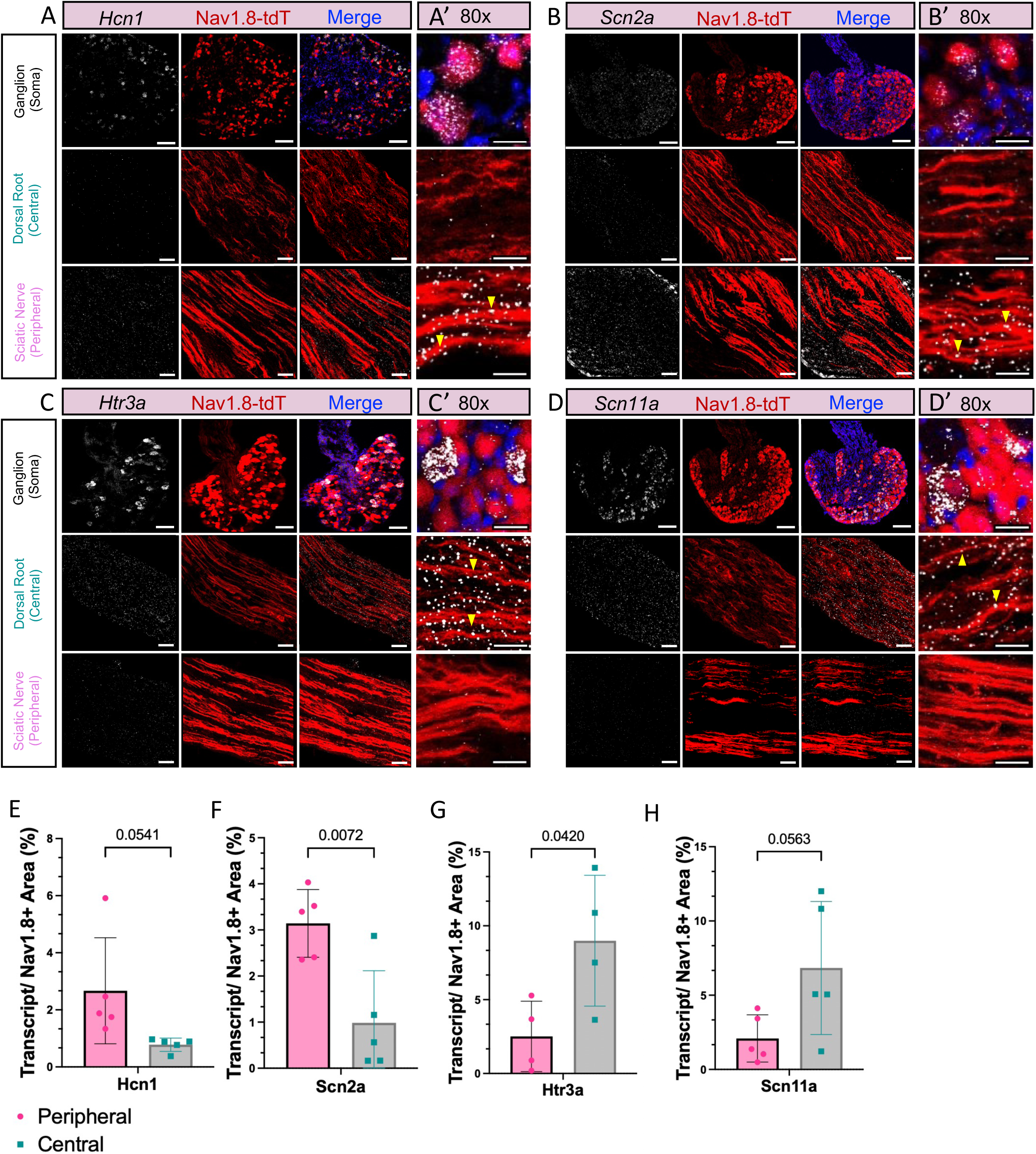
Transcripts encoding locally translated ion channels have axon-specific expression. A-D) Representative 20x fluorescent images of *Hcn1* (A)*, Scn2a* (B), *Htr3a* (D), *and Scn11a* (D) mRNA puncta (visualized by RNAScope *in situ* hybridization) and TdTomato-labeled Nav1.8+ (Nav1.8-tdT) neurons in DRGs, dorsal roots and sciatic nerves, representing DRG soma, central axons, and peripheral axons, respectively. Scale bar = 100 μm (soma) and 40 μm (axons). A’-D’) Higher magnification images demonstrating the location and abundance of *Hcn1* (A’)*, Scn2a* (B’), *Htr3a* (D’), *and Scn11a* (D’) mRNA puncta within Nav1.8-tdT+ soma in the ganglia, and axon in the dorsal root and sciatic nerve. Scale bar = 10 μm. E-H) Quantification of the percent area occupied by *Hcn1* (E), *Scn2a* (F), *Htr3a* (G), and *Scn11a* (H) mRNA puncta within Nav1.8-tdT+ axons in dorsal roots and sciatic nerves. Statistical significance determined by two-way ANOVA with Tukey’s multiple comparisons test and multiple unpaired t-tests (p-value ≤ 0.05). Data are represented as mean ± SEM. n=1-2 sciatic nerves samples/mouse and 1-3 dorsal roots samples/mouse from 4-5 mice.

### RBPs are positioned to orchestrate the subcellular localization of axonal translatomes

Our TRAP-Seq, PLA and RNAscope findings indicate that many of the channels and receptors critical for initiating and propagating somatosensation are locally produced within either the central or peripheral axon, and that the corresponding mRNAs are highly localized to one or the other axon as well. An important step in enabling such highly localized and regulated translation is the sorting and transport of mRNAs to distinct subcellular regions. Sorting of mRNAs to specific locations often depends on structural elements within mRNA sequences; axonal mRNA sorting is usually coordinated through RNA binding protein (RBP) recognition of select localization motifs within the 3’ untranslated region (3’UTR) of the target mRNA ^34^. RBP-bound mRNAs assemble into ribonucleoprotein (RNP) transport granules, characterized by their ability to undergo liquid-liquid phase separation and be transported directly or indirectly along microtubules by motor proteins ^35–37^.

To investigate the potential logic for differential RNA sorting to central and peripheral axons, we bioinformatically analyzed the 3’UTRs of axonally translated mRNAs, with the goal of identifying RBPs that selectively bind and traffic mRNAs to one or the other axon ^38^. RBPs were identified as candidates for mRNA sorting if they fulfilled two criteria: 1) multiple binding motifs for that RPB are in the 3’UTR of key axonally translated mRNAs and 2) the RBP is predicted and/or demonstrated to undergo phase separation and be present in RNA granules ^39–41^. A heatmap displaying the number of individual motifs per 3’UTR was used to rank RBPs that were significantly more associated with peripherally or centrally translated mRNAs (Fig. S6A). We determined that 16 RBPs similarly bound the 3’UTRs of peripherally and centrally translated mRNAs, 84 RBPs had more binding sites within peripherally translated mRNA transcripts, and 24 RBPs had more binding sites within centrally translated mRNA transcripts (Fig. S6A).

We further investigated the three cohorts of RBPs using the STRING database protein-protein interaction functional annotation analysis (Fig. S6B-D) ^42^. As predicted, all cohorts were enriched for RBPs involved in RNP granules cellular components. The set of RBPs that are predicted to bind equally to mRNAs present in the peripheral and central axons were enriched for nuclear proteins (Fig. S6B). Interestingly, peripheral-associated RBPs were instead enriched for proteins with specific neuronal functions in the synapse, dendrite, and distal axon (Fig. S6C). RBPs predicted to bind to the centrally located and centrally translated mRNA were enriched for cytoplasmic proteins that are involved in both RNP and stress granules (Fig. S6D). These data prompted us to examine the possibility that some of these RBPs may be constituents of cytoplasmic RNP granules involved in selective axonal transport and translation.

If individual RBPs are implicated in mRNA sorting and axon-specific translation, we would expect to see spatially selective expression of these RBPs in the axons *in vivo*. Therefore, several candidate RBPs were further investigated to determine if *in vivo* axonal localization patterns correlate with their predicted mRNA binding patterns. We focused on the localization patterns of Splicing factor proline- and glutamine-rich (SFPQ), Fus-interacting protein 1/ serine and arginine rich splicing factor 10 (Fusip1/SRSF10), and Embryonic lethal, abnormal vision, drosophila-like 1 (HuR/ELAVL1). Interestingly, SFPQ contains 3’UTR binding sites on multiple peripherally translated mRNAs required for axonal and synaptic function including *Hcn1, Gabrb3, Calb1,* and *Dnajc6* (Fig. S6A). In contrast, SRSF10 contains 3’UTR binding sites on multiple centrally translated mRNAs encoding ion channels and receptors related to pain signaling: *Htr3a, Htr5a, Trpv1, P2ry2* and *Chrm1* (Fig. S6A). ELAVL1 has predicted binding sites on almost all locally translated mRNAs regardless of axon specificity (Fig. S6A).

Fluorescence immunohistochemistry paired with particle analysis demonstrates that SFPQ protein is enriched in peripheral axons of the sciatic nerve (Fig. 6A,D), consistent with our prediction that SFPQ preferentially binds to mRNAs in the peripheral axons. In contrast, we find that SRSF10 protein is equally present in both central axons of the dorsal root and peripheral axons of the sciatic nerve (Fig. 6B,D). The third RBP tested, ELAVL1, is present in both the dorsal root and the sciatic nerve but is largely restricted to the nuclei of glia and other supporting cells (Fig. 6C,D). Thus, both SFPQ and SRSF10 are appropriately localized to contribute to trafficking and translation of mRNAs in axons, and SFPQ is appropriately located to promote selective sorting towards the peripheral axon. Our data also indicate another difference between the RBP puncta in the central versus peripheral axons: the puncta of SFPQ and SRSF10 were consistently larger within the sciatic nerve than in the dorsal root. This difference in size of the RBP-puncta suggest that there may be distinct features of RNP granules in each axon (Fig. 6E).

**Figure 6.**
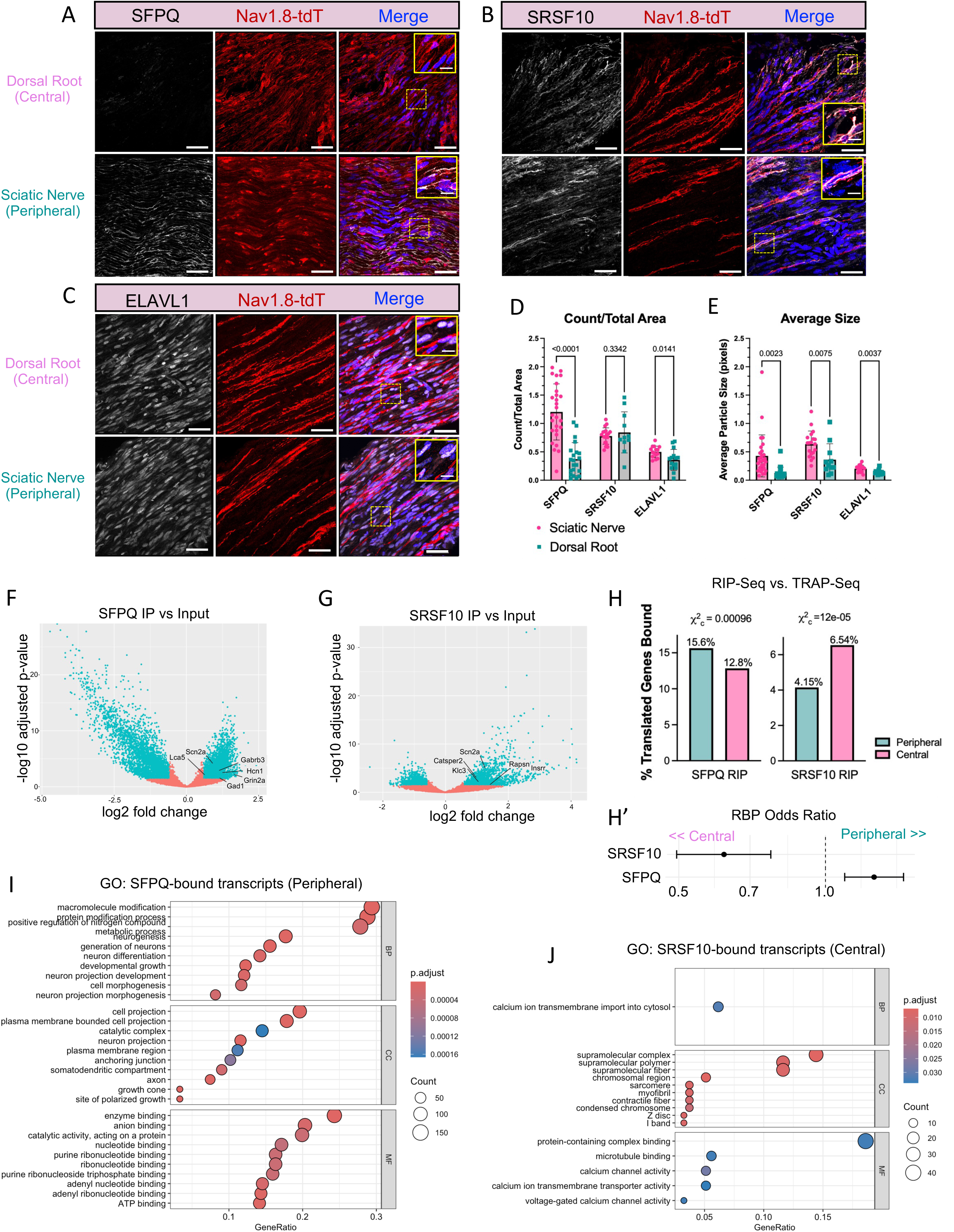
Select RBPs orchestrate axon-specific mRNA transport. A-C) Representative 60x fluorescent immunohistochemistry images of SFPQ (A), SRSF10 (B), and ELAVL1 (C) protein in TdTomato-labeled Nav1.8+ (Nav1.8-tdT) axons in dorsal roots and sciatic nerves, representing DRG central and peripheral axons, respectively. The solid yellow box shows a 4x magnified image of the area surrounded by yellow dotted lines, demonstrating the location and abundance of protein within axons and nuclei of supporting cells. Scale bars: Full image = 40 μm; inlaid image = 10 μm. D, E) Quantification of the number (D) and size (E) of SFPQ, SRSF10, and ELAVL1-positive puncta within Nav1.8-tdT+ axons in dorsal roots and sciatic nerves. Statistical significance determined by two-way ANOVA with Tukey’s multiple comparisons test. Data are represented as mean ± SEM. (n=1-2 sciatic nerves samples/mouse and 1-3 dorsal roots samples/mouse from 4-5 mice). F, G) Volcano plots from RIP-Seq experiments demonstrating the mRNA transcripts with significantly higher expression in SFPQ IP (F) and SRSF10 IP (G) samples as compared to the input fraction from the same DRG neuron cell culture experiments. Select locally translated mRNAs bound by SFPQ and SRSF10 are indicated on the volcano plots. (n= 4 sets; p.adj ≤ 0.05; fold-change ≥ 1). Chi-square analysis comparing RBP-bound transcripts to compartment-resolved TRAP translatomes. Contingency tables were generated by intersecting RIP-seq–identified SFPQ-bound or SRSF10-bound mRNAs with the background-filtered peripheral and central axon gene sets. Chi-square testing revealed a significant association between RBP identity and axonal compartment (p ≤ 0.0001), with SFPQ-bound transcripts enriched among peripherally translated mRNAs and SRSF10-bound transcripts enriched among centrally translated mRNAs. H’) Odds-ratio analysis quantifying the direction and magnitude of these biases. SFPQ-bound transcripts were 1.26-fold more likely to belong to the peripheral axon translatome, whereas SRSF10-bound transcripts were 0.38-fold less likely to be peripheral, indicating preferential association with centrally translated mRNAs. I, J) Gene ontology (GO) enrichment analysis visualized as a dot plot, where each dot represents the most significantly enriched biological processes (BP), cellular compartments (CC), or molecular functions (MF) from the SFPQ-associated peripheral axon regulon (I) and SRSF10-associated central axon regulon (J) (p.adj ≤ 0.05).

### Select RBPs guide the axonal sorting and transport of functional RNA regulons

To determine whether SRSF10 and SFPQ bind and sort mRNAs for axonal translation, we performed RNA Immunoprecipitation paired with sequencing (RIP-Seq) in DRG neurons. In our experiments, SRSF10 and SFPQ protein were immunoprecipitated from DRG neurons grown in culture for 5 days, and mRNAs from these immunoprecipitates and from rabbit IgG negative controls were extracted and sequenced (Fig. S7A-D, Table S13). We identified the mRNAs that are significantly enriched in the immunoprecipitates compared to the inputs. The mRNAs enriched in the SFPQ and SRSF10 immunoprecipitates include transcripts encoding several ion channels and receptors that are locally translated within axons, including *Hcn1*, *Scn2a*, *Scn11a*, and *Catsper2* (Fig. 6F,G; Tables S14,15).

To define the functional identity of each RBP’s cargo, we first performed GO enrichment analysis on the full set of SFPQ- and SRSF10-bound transcripts identified by RIP-seq, independent of whether these RNAs appeared in the local translatomes (Fig. S7E-F’; Tables S16,17). SFPQ-bound mRNAs were enriched for biological processes related to neuronal projection organization and assembly, including axon shaft components, distal axons, growth cones, and synaptic structures, consistent with SFPQ’s established role in long-range axonal RNA trafficking (Fig. S7E; Table S16). In contrast, SRSF10-bound mRNAs were enriched for pathways related to mRNA processing and RNA splicing, but also contained transcripts encoding multiple structural and signaling components found in central axons—including presynaptic active zone proteins, calcium channel complexes, and microtubule motor complexes (Fig. S7F,F’; Table S17). These results indicate that SFPQ and SRSF10 engage functionally distinct RNA regulons, each composed of transcripts that support different aspects of neuronal physiology.

Having established the global functional signatures of these RBP-bound regulons, we next asked whether SFPQ and SRSF10 preferentially associate with mRNAs that are locally translated in peripheral versus central axons. Chi-square analysis comparing RIP-seq targets to the TRAP datasets revealed a significant association between RBP identity and axonal compartment (Fig. 6H). SFPQ-bound transcripts were selectively enriched among peripherally translated mRNAs, whereas SRSF10-bound transcripts were disproportionately enriched among centrally translated mRNAs. Odds-ratio analysis further quantified these biases, showing that SFPQ-bound transcripts were 26% more likely to be peripheral, while SRSF10-bound transcripts were 38% less likely to be peripheral (i.e., 62% more likely to be central) (Fig. 6H′). These findings demonstrate that each RBP binds distinct RNA regulons that preferentially traffick to one or the other axon of sensory neurons.

To further resolve the biological functions of these compartment-directed regulons, we performed GO enrichment analyses on SFPQ- and SRSF10-bound transcripts that overlapped with the peripheral and central axonal translatomes, respectively (Fig. 6I,J). The SFPQ-associated peripheral translatome was enriched for biological processes related to metabolic pathways, axon development, neuron projection morphogenesis, as well as molecular functions linked to ATP binding and enzymatic activity. These categories align with the metabolic and structural demands of the distal sensory terminal and suggest that SFPQ coordinates a peripheral regulon tuned for axon maintenance, remodeling, and localized metabolism (Fig. 6I). In contrast, the SRSF10-associated central translatome was enriched for pathways governing calcium ion transmembrane import and cytoskeletal organization, and molecular functions such as microtubule binding and voltage-gated calcium channel activity (Fig. 6J). These functions reflect the specialized physiology of central presynaptic terminals, where precise control of calcium influx, cytoskeletal scaffolding, and synaptic machinery is essential for neurotransmitter release. Together, these results reveal that SFPQ and SRSF10 enable polarized RNA regulons that contribute to the divergent structural and synaptic identities of peripheral and central DRG axons. By coordinating the selective transport and local translation of distinct gene cohorts into each axonal branch, these RBPs provide a mechanistic framework for how a single sensory neuron maintains functional asymmetry across its long expanse.

## DISCUSSION

Spatial translatomics and high-resolution microscopy have revolutionized our ability to identify locally translated mRNAs and understand both the underlying mechanisms and the functional consequences of these subcellular specializations. Here we show that neuronal axon projections from the same cell can have distinct translatomes to fulfill specialized functional requirements. While molecules that modulate synaptic strength and nociceptive signaling are locally translated in central axons of the somatosensory neurons, peripheral axons invest in sustaining the structural and metabolic demands of long-range projections and regeneration. Our studies also reveal the logic behind axonal sorting of select mRNAs; the RBPs, SFPQ and SRSF10, preferentially bind the 3’UTRs of mRNAs that are axonally translated, with SFPQ implicated in sorting to peripheral axons and SRSF10 emerging as a regulator of centrally translated mRNAs. Furthermore, the set of mRNAs that are locally translated in the peripheral and central axons include multiple ion channels and receptors related to neuronal firing, nociceptive processing, and regenerative capacity of sensory neurons. Thus, regulated translation has the potential of selectively and dynamically modifying signal transmission within individual axon branches both in normal physiology and in pathophysiology.

Perhaps surprisingly, our studies indicate that many mRNAs that are translated within axons encode plasma membrane-bound ion channels and receptors. It is therefore appropriate to question whether axons are capable of the translational and secretory pathways required for membrane targeting and integration. Early electron microscopy studies did not detect ribosomes, rough endoplasmic reticulum, or Golgi apparatus in axons ^43^. More recently, monosomes, small diameter ER tubules, and Golgi satellites have been identified in peripheral axons ^16,44–48^. It has been suggested that axonal ion channels can be shuttled from axonal Golgi satellites to the plasma membrane through Rab6-positive exocytic vesicles ^49^. In addition, atypical membrane proteins that are core-glycosylated rather than N-glycosylated may bypass the Golgi apparatus and be more directly shuttled to the axonal plasma membrane ^50,51^. Indeed, both the peripheral and central axonal translatomes include glycosylating enzymes (*Galnt5* in peripheral axon dataset and *St6galnac2* in central axon dataset). Several key locally translated mRNAs identified here, including *Scn2a, Gabrb3*, and *Grin2a,* encode proteins that have been observed to be core-glycosylated in axonal membranes ^50^. Taken together, specialized axonal translation, processing, and secretion of membrane bound ion channels and receptors is both feasible and likely in distant axons.

The discovery that DRG central axons locally translate transcripts encoding ion channels, neurotransmitter receptors, and synaptic scaffold proteins suggests that presynaptic terminals in the spinal cord possess an intrinsic capacity for molecular remodeling. Local synthesis of glutamatergic and GABAergic receptor components (*Grin1/2, Grm1/3/5, Gabra1–5*), along with scaffolding proteins such as *Nrxn1, Nlgn1*, and *Dlg2/3*, could permit rapid, activity-dependent adjustment of receptor composition and synaptic strength within the dorsal horn. Similarly, translation of voltage-gated calcium channel subunits and presynaptic release machinery (*Stx1b, Vamp2*) provides a mechanism to tune excitability and neurotransmitter release in response to somatosensory stimulation. Together, these findings indicate that DRG central axons are active, translationally competent structures capable of locally modifying synaptic plasticity, thereby contributing to the habituation and/or sensitization of nociceptive circuits.

Our findings reveal that the neuromodulatory serotonin receptor, 5-HTR3A, is selectively located and translated in central axons, indicating that serotonergic signals in the spinal cord directly modulate primary somatosensory inputs. Indeed, axons from serotonergic neurons of the raphe nuclei are known to descend to the dorsal horn of the spinal cord, and ∼15% of axo-axonic contacts with DRGs are serotonergic ^52,53^. While previous models emphasized indirect modulation of somatosensory neurons through GABAergic interneurons that express excitatory 5-HTR3A receptors ^54,55^, our findings suggest serotonergic stimulation of the central terminals of primary afferents themselves. Therefore, adjustments of pain signaling and sensation within the excitatory, glutamatergic central axon terminals may provide an important initial site of pain modulation. Together, these results support a model in which the local translation of 5-HTR3A equips DRG central axons to integrate descending signals with ongoing sensory activity, shaping early stages of nociceptive information processing ^53,56^.

Local translation also provides a dynamic mechanism for structural remodeling within axonal compartments. In central DRG axons, the local synthesis of cytoskeletal and scaffolding proteins supports the dynamic modification of presynaptic architecture and receptor alignment, enabling synaptic plasticity within the dorsal horn. In peripheral DRG axons, local translation is poised to support the structural remodeling required for axonal maintenance, growth-cone dynamics, and interactions with the extracellular environment. The capacity for local synthesis of cytoskeletal and translational components may underlie the unique regenerative potential of peripheral, but not central, axons. Peripheral nerve regeneration depends on both neuron-intrinsic components and signaling pathways initiated by glial and immune cells in the local microenvironment ^57,58^. Dynamic remodeling of the axon cytoskeleton and functional connections to cell-extrinsic ECM proteins and glial cells may allow damaged sensory fibers to restore axonal integrity, rebuild growth cones, and re-establish functional connections after injury.

Furthermore, local synthesis of ribosomal components and other protein-synthesis machinery suggests that peripheral axons can replenish or repair their translational apparatus on site, supporting the high biosynthetic demands of regeneration after mechanical injuries or other cell stresses^59^. Together localized regulation and synthesis of cytoskeletal outgrowth and translational machinery provide peripheral axons with regenerative capacity in response to nerve injury.

Our data indicate that local translation also can play a role in modulating axonal physiology by dynamically adjusting the complement of ion channels along DRG neurons. Although differential ion channel insertion in axonal membranes has long been known to influence neuroplasticity by altering membrane composition and excitability ^60–62^, our findings reveal that genes encoding several voltage-gated sodium channels are locally and differentially translated in the axons. There are nine distinct genes that encode different voltage-gated sodium channels, Nav1.1–1.9 ^63^, and individual neurons of the CNS and PNS each express a subset of these channel genes ^32,64,65^. Our data indicate that *Scn2a*, which encodes Nav1.2, is translated in peripheral axons. Nav1.2 is a tetrodotoxin-sensitive (TTX) channel with fast kinetics ^32,64–67^. In addition, we find that the Hyperpolarization Activated Cyclic Nucleotide Gated Potassium Channel 1 (HCN1) is also selectively translated in the peripheral axons. HCN1 exhibits rapid kinetics, enhances action potential initiation and enables increased action potential frequency ^68–70^.

Peripheral axons resemble dendrites in that they are the site of initial stimulation and signal integration^71^ and we hypothesize that HCN1 and Nav1.2 may enable initiation and propagation of action potentials in both CNS dendrites and DRG peripheral axons. In contrast, we find that *Scn11a*, which encodes Nav1.9, is preferentially translated in the central axons. Unlike Nav1.2, Nav1.9 is a slowly activating/ inactivating, TTX-resistant channel that amplifies subthreshold signals ^32,64–67^ Nav1.9 is predominantly expressed in nociceptive neurons in the PNS, and we suggest that this channel may amplify subthreshold pain signals in the central axon and so adjust the electrophysiologic input to the spinal cord^32^. Mutations or altered expression of many of these newly identified axonally translated components have been implicated in human pain syndromes. Gain-of-function mutations of Scn11a, a component of the central axon translatome, results in episodic cold pain sensation and painful peripheral neuropathy in people, while loss of function mutations disrupt inflammatory and cold-triggered pain sensation in mice and humans ^32,72,73^. Similarly, null mutations of *Hcn,* which is translated in peripheral axons, result in reduced neuropathic pain and cold allodynia after nerve injury, when compared to wild-type mice after nerve injury^74^. Therefore, precise control over *Scn11a* and *Hcn1* localization and translation is likely required for appropriate pain sensation and may play a prominent role in neuropathic pain syndromes ^72,73^.

A common and often devastating cause of neuropathic pain is chemotherapy-induced peripheral neuropathy. Here we integrate the analysis of locally translated mRNAs in DRG soma and axons with the genesets that change following paclitaxel treatment, a chemotherapeutic agent known to cause axonal degeneration and peripheral neuropathy. We found that paclitaxel alters expression of many mRNAs encoding proteins critical for axonal transport and vesicle localization, and these mRNAs are locally translated in all of the subcellular domains, indicating that intracellular transport is broadly impacted by paclitaxel. In addition, mRNAs with altered expression in response to paclitaxel include many mRNAs translated in the peripheral axons that contribute to cytoplasmic translation, neuronal projection extension, and environmental sensing, whereas paclitaxel-altered transcripts that are translated in central axons modulate synaptic and secretory pathways. These findings reveal that neuropathic injury elicits widespread and domain-specific adaptations in protein translation that may collectively contribute to chronic pain phenotypes.

A key question addressed here is how specific RNA regulons, sets of functionally related mRNAs with synchronized expression, are coordinately transported and sorted to a specific axon. RBPs are known to regulate RNA regulons at the level of splicing, nuclear export, transport, localization, and stability ^75^. It has been shown that co-regulation of RNA regulons by specific RBPs is required for axon viability ^30,76^. In this study, we implicate two RBPs in the organization of central and peripheral axonal translatomes. SFPQ and SRSF10 were initially identified as RNA splicing factors containing both RNA recognition motifs (RRMs) and intrinsically disordered regions ^77–79^. In addition to its role in splicing, SFPQ has been identified as a component of axonal RNP granules and enables axonal transport of mRNA cargoes ^30,80–82^. While less is known about SRSF10, it has been shown to regulate metabolic processes such as glycolysis ^83^, and has also been functionally linked to neurogenesis and spermatogenesis ^84–86^. Here we show that SFPQ and SRSF10 bind multiple locally translated mRNAs and that the subcellular distribution of these two RBPS is strikingly different, with SFPQ enriched in peripheral axons and SRSF10 present in both axons. RIP-Seq and GO analyses demonstrate that the SFPQ-associated regulon was enriched for metabolic pathways, neuronal projection morphogenesis, and distal axon maintenance, while the SRSF10-associated regulon was enriched for calcium channel complexes, cytoskeletal scaffolding, and presynaptic active zone components. These patterns support a model in which polarized RNA regulons reinforce the divergent structural and synaptic identities of peripheral and central axons.

While many RBPs haven been implicated in axonal trafficking, our data indicate that SFPQ is involved in selectively sorting mRNAs to the peripheral, rather than the central axons. This novel role joins a host of cytoplasmic functions qualifying SFPQ as a master regulator of axonal survival^80^. For example, in peripheral neurons, SFPQ directly binds a kinesin complex containing the motor, KIF5A, and adaptor, KLC1, to transport mRNAs encoding axonal survival factors, such as *Laminb2* and *Bclw*^29,30,87^. Indeed, both SFPQ knockdown and selective interruption in transport activity led to axon degeneration in sensory neurons. Moreover, cytoplasmic aggregation of SFPQ is implicated in several neurodegenerative disorders including, frontotemporal lobar degeneration (FTLD), and Alzheimer’s disease, as well as sensory neuropathies including chemotherapy-induced peripheral neuropathy, while SFPQ mutations can cause motor neuron dysfunction and amyotrophic lateral sclerosis (ALS) ^80–82^. Our findings now extend our understanding of SFPQ and degeneration, with evidence that SFPQ-dependent RNA granule transport and sorting is likely to contribute to loss of epidermal innervation by peripheral somatosensory axons in neuropathy.

SRSF10 has known roles in regulating RNAs involved in neurogenesis and myelination, but has not previously been implicated in axonal trafficking or neurodegeneration. While SFSF10 is present at similar levels in both the central and peripheral axons, our data suggest that it has a preferential role in transcripts targeted to the central axons, including transcripts encoding calcium channels and regulators. Interestingly, conditional knockouts of SRSF10 in neural progenitor cells results in impaired performance on multiple learning and memory tasks, which involve localized calcium regulation ^88^.

The comprehensive comparisons of central and peripheral axon translatomes in DRG neurons assembled here provide critical resources for understanding the basis of compartmentalized neural function. The axonal translatomes reveal that synaptic, electrophysiologic, structural, and regenerative properties of neurons can be rapidly and locally modulated by changes in translation in response to environmental cues. Identification of distinct RBPs that coordinate and guide intracellular transport and translation of RNA regulons reveals key components that determine the asymmetry of neuronal projections required for functional circuitry across distinct biological spheres. Overall, the spatial regulation of translation, and the selective routing of RNA regulons by SFPQ and SRSF10, emerges as a unifying mechanism that preserves neuronal function and adaptability across the diverse environments encountered by sensory axons, and helps explain how neuropathic stress reshapes molecular identity across subcellular compartments.

## RESOURCE AVAILABILITY

### Lead Contact

- Requests for further information and resources should be directed to the lead contact, Rosalind Segal.

### Materials Availability

- This study did not generate new unique reagents.

### Data and code availability

- TRAP-seq and RIP-seq data have been deposited at GEO:GSE289550 and GEO:GSE289551 and will be publicly available as of the date of publication.
- TRAP-seq data are also available for browsing and analysis via the Pain-seq multi-omic data resource at http://painseq.shinyapps.io/CompartmentTRAP/.
- Novel code used to generate the web resource has been deposited to GitHub (http://github.com/Renthal-Lab/CompartmentTRAPShiny).
- Any additional information required to reanalyze the data reported in this paper is available from the lead contact upon request.

## Supporting information

TRAP Supplemental Tables

RIP Supplemental Tables

## ACKNOWLEDGEMENTS

RNA Sequencing was performed in the University of California-San Francisco Genomics CoLab by Michael Adkisson, Andrew Schroeder, Andrea Barczak, and Walter Eckalbar. RIP Sequencing was performed by the MBCF: Genomics Core Facility at Dana-Farber Cancer Institute by Zack Herbert and Maura Berkeley. This work was supported by NIH grants R01-NS050674 and R01-CA205255 (RAS) and T32-AG000222 (ES). We thank David Ginty, Mike Greenberg, and Charles D Stiles for helpful comments on the manuscript, and Myriam Heiman, Sarah Pease-Raissi, and Ozge Tasdemir-Yilmaz for help with TRAP-Seq methods.

## AUTHOR CONTRIBUTIONS

ESS conceptualized and designed the study, performed the experiments, analyzed the data, and wrote the manuscript. EN assisted with tissue processing, performed immunohistochemical experiments, ran bioinformatic analysis, and reviewed and edited the manuscript. MFP provided technical expertise, maintained mouse colonies, and reviewed and edited the manuscript. JZP assisted with tissue processing, conducted immunohistochemical experiments, and review and edited the manuscript. SAB and WR developed the web resource and performed bioinformatic analysis. RAS conceptualized and designed the study, supervised the research, contributed to the data interpretation, and provided critical revisions to the manuscript.

## STAR METHODS

## EXPERIMENTAL MODEL AND STUDY PARTICIPANT DETAILS

### Mouse Lines and Animal Care

All experimental procedures were conducted in accordance with the National Institutes of Health (NIH) guidelines and were approved by the Dana-Farber Cancer Institute (DFCI) Institutional Animal Care and Use Committee (IACUC).

*Mouse Strains:* L10a-eGFP^fl/fl^ (B6;129S4-*Gt(ROSA)26Sor^tm9(EGFP/Rpl10a)Amc^*/J) mice were the generous gift of Myriam Heiman (Massachusetts Institute of Technology) ^26^. Nav1.8^cre^ (B6.129-Scn10a^tm2(cre)Jwo^/H) and TdTomato^fl/fl^ (B6.Cg-*Gt(ROSA)26Sor^tm14(CAG-tdTomato)Hze^*/J) mice were kindly provided by David Ginty (Harvard Medical School). For *in vivo* TRAP studies, Nav1.8^cre/+^ and L10a-eGFP^fl/fl^ mice were crossed to generate Nav1.8^cre^;L10a-eGFP^fl/fl^ mice (GFP+) and L10a-eGFP^fl/fl^ (GFP-) littermate controls. For tissue collection and histology, Nav1.8^cre^ and TdTomato^fl/fl^ mice were crossed to generate Nav1.8^cre^;TdTomato^fl/fl^ mice with labelled sensory neurons.

Mice were group housed in a temperature-controlled environment (22 ± 2°C) with a 12-hour light/dark cycle and had free access to food and water. Mice aged P3-5 were used for all experiments. For TRAP experiments, 13-17 littermate animals of both sexes were pooled by genotype (GFP positive or negative) and used to generate TRAP samples. Pooling of both sexes was required to generate enough tissue material for TRAP-Sequencing. For tissue collection and histology, all TdTomato-expressing littermate animals of both sexes were used to generate tissue samples for downstream applications.

### Primary Rat Cell Cultures

Timed pregnant Sprague-Dawley rats were purchased from Charles River for primary DRG neuron isolation. Embryos from timed pregnant Sprague-Dawley rats (aged E15) were used for primary cell culture isolation. All embryos, presumably of both sexes, were used. DRGs were dissected, dissociated, and plated in Matrigel (1:45; Thermo Fisher Scientific)-coated p35 dishes. DRG cultures were maintained in NeuroBasal medium supplemented with 2% B27, 1% Glutamax, 1% penicillin and streptomycin, 0.08% glucose, 1–100 ng/ml nerve growth factor (NGF)/brain-derived neurotrophic factor (BDNF; PeproTech), and 0.5 µM Cytarabine (AraC). Cultures were maintained in incubators at 37°C with 7.5% CO_2_ for 5 days.

## METHOD DETAILS

### Translating Ribosomal Affinity Purification (TRAP)

TRAP protocol was adapted from Myriam Heiman ^26^. Roughly 15 Nav1.8-Cre;L10a-eGFP mice aged P3-5 and L10a-eGP littermate controls were euthanized by CO_2_ inhalation and cervical dislocation. Lumbar (L1-6) dorsal root ganglia (Soma), sciatic nerve (PA), and lumbar dorsal roots and spinal cords (CA) were quickly manually dissected in Dissection Buffer (1X HBSS, 2.5mM HEPES-KOH [pH7.4], 25mM Glucose, 4mM NaHCO_3_, 100µg/mL cycloheximide) from pooled Soma and CA compartment tissues were immediately homogenized in ice-cold Polysome Extraction Buffer (20mH M HEPES-KOH [pH 7.4], 5mM MgCl_2_, 150 mM KCl, 0.5mM DTT, 100µg/mL cycloheximide, protease inhibitors (EDTA-free), 40 U/mL RNAsin, 20 U/mL Superasin) using glass Dounce homogenizers. Pooled PA compartment tissue was enzymatically digested (0.1% Collagenase II, 0.2% Hyaluronidase, 40 U/mL RNAsin, 20 U/mL Superasin) in Dissection Buffer for 5 min at room temperature, then resuspended in ice-cold Polysome Extraction Buffer and Dounce homogenized. Homogenates were centrifuged for 10 minutes at 2,000 × *g*, 4 °C, to pellet large cell debris, and 1% NP-40 (AG Scientific, San Diego, CA) and 30 mM DHPC (Avanti Polar Lipids, Alabaster, AL) were added to the supernatant. After incubation on ice for 5 minutes, the lysates were centrifuged for 10 minutes at 20,000 × *g*, 4 °C. Supernatants were pre-cleared with unbound Pierce Protein L Magnetic Beads at 4 °C for 1 hour. Custom anti-GFP antibody (Htz-GFP-19C8; RRID:AB_2716737 and Htz-GFP-19F7; RRID:AB_2716736) coated Pierce Protein L Magnetic Beads were added to the supernatant and incubated at 4 °C with end-to-end rotation overnight. Beads were subsequently collected on a magnetic rack, washed three times with high-salt IP Wash Buffer (20 mM HEPES [pH 7.4], 350 mM KCl, 5 mM MgCl_2_, 1% NP-40, 0.5mM dithiothreitol, 100 μg/ml cycloheximide, 3% bovine serum albumin [IgG Protease-free]) and immediately placed in Lysis Buffer with β-ME (Absolutely RNA Nanoprep Kit) (Agilent Technologies, Santa Clara, CA) and incubated for 10 mins at room temperature. Eluted RNA in Lysis Buffer was removed from beads on a magnetic rack. RNA was further purified using the Absolutely RNA Nanoprep Kit with column purification and DNase digestion. RNA integrity and concentration was measured on an Agilent 2100 Bioanalyzer System with RNA 6000 Pico Kit (Agilent Technologies, Santa Clara, CA); amounts of RNA were estimated as <100pg for axon samples and <1ng for soma samples. Four biological replicates were performed for CA and PA tissues, and eight replicates were performed for somatic tissues.

### Low Input Total RNA Sequencing and Analysis

Low input total RNA Sequencing (>10pg total RNA or 1-500 cells) was performed at the University of California-San Francisco Functional Genomics Core Facility. RNA libraries were prepared using the SMART-Seq v4 Mouse Kit (Takara Bio Inc.) and sequenced on an Illumina HiSeq 4000 instrument via 50bp sequencing. Sequences were aligned to mouse reference genome with STAR_2.7.2b ^27^. RNA-seq analysis of raw counts was performed in R v4.3.2 for sample-level quality control (principal component analysis), Differential Gene Expression testing for pairwise, group comparisons (DESeq2) ^31^, and Functional Annotation (enrichGO) ^89,90^.

### Integrative Cross Dataset Analysis

To identify genes enriched in paclitaxel-treated cells relative to naïve cells within each cell type, we performed differential gene expression analysis using Seurat’s FindAllMarkers function with default settings on data generated by Renthal and colleagues^29,30^. P-values were subsequently adjusted for multiple comparisons using the Benjamini-Hochberg procedure (FDR). We then intersected the differentially expressed genes from each cell subtype with those identified in our TRAP-seq data across anatomical compartments. For each resulting intersected gene list, we conducted Gene Ontology (GO) enrichment analysis – covering biological process, molecular function, and cellular component categories – using the normalized gene counts from our TRAP-seq dataset serving as the background set.

### Puromycin Injection

Nav1.8^cre^;TdTomato^fl/fl^ mice were injected with Puromycin Dihydrochloride (#P33020) (Research Products International, Mount Prospect, IL) at a concentration of 225mg/kg (0.9mg/mouse) delivered via intraperitoneal injection (30 μL in dH2O). After 25 min, mice were euthanized by CO_2_ inhalation followed by cervical dislocation.

### Mouse Histological Methods

Dorsal root ganglia (Soma), sciatic nerves (PA), and dorsal roots (CA) were rapidly dissected from euthanized mice in ice-cold 1X PBS and placed immediately fixed in 4% PFA overnight. Tissues were washed 3×5min with 1XPBS, then prepared for embedding with 10% to 30% sucrose gradient over 3 days, embedded in NEG-50 medium (VWR, Radnor, Pennsylvania), and sectioned at 16μm on a cryostat.

### Proximity Ligation Assay

Proximity ligation was performed according to the Duolink In Situ Proximity Ligation (Millipore Sigma, Burlington, MA) protocol. In brief, 20μg of primary anti-Puromycin (Sigma, #MABE343; RRID:AB_2566826), anti-5HTR3A (Thermo Scientific, #PA5-77746; RRID:AB_2735937), anti-SCN11A (Alomone, #ASC-017; RRID:AB_2040200), anti-SCN2A (Alomone, #ASC-002; RRID:AB_2040005), and anti-HCN1(Alomone, #APC-056; RRID:AB_2039900) antibodies were conjugated to Duolink probes using the Duolink In Situ Probemaker Kit (Millipore Sigma, Burlington, MA). Frozen tissue sections were accommodated to room temperature for 30 mins. Slides were then permeabilized with 0.1% PBS-Triton-X-100 (PBST) and blocked with Duolink Blocking Solution. Conjugated primary antibodies were added to slides and incubated at 4 °C overnight. Duolink ligation and polymerization steps were performed to amplify fluorescent signal, followed by two wash steps. Coverslips were mounted with ProLong Gold Antifade Mountant complete with DAPI (ThermoFisher).

### RNAscope In Situ Hybridization

In situ hybridization was performed according to the Co-detection Immunohistochemistry with RNAscope Fluorescent Multiplex Reagent (ACDBio, Burlington, MA) protocol for fixed-frozen tissue samples. In brief, sample slides were accommodated to room temperature for 30 mins. Sections were then prepared by washing with 1X PBS before baking for 30 mins at 60 °C. Sections were then post-fixed with 4% PFA for 15 minutes at 4 °C. Samples were dehydrated through serial incubations in 50%, 70%, and 100% Ethanol. Samples were treated with hydrogen peroxide for 10 min and then underwent antigen retrieval with RNAscope Target Retrieval Solution in a steamer basket for 5 minutes. Primary anti-RFP antibody (VWR, #RL600-401-379) was applied to slides and incubated at 4 °C overnight. Samples then underwent post-primary fixation in 4% PFA for 30 mins, followed by Protease Plus incubation for 30 mins in a HybEZ Oven at 40 °C, probe hybridization for 30 mins at 40 °C, AMP hybridization for 30 mins at 40 °C, HRP signal development for 15 mins at 40 °C, Opal 690 Reagent (Akoya Biosciences) incubation for 30 mins at 40 °C, and HRP blocking for 15 mins at 40 °C. Samples were then incubated with secondary antibody (Invitrogen: AlexaFluor H+L) for 60 mins, followed by two wash steps. Coverslips were mounted with ProLong Gold Antifade Mountant complete with DAPI (ThermoFisher).

### Fluorescent Immunohistochemistry

Fixed frozen sample slides were accommodated to room temperature for 30 mins. Samples were then permeabilized 3 x 5 min with 0.1% PBS-Triton-X-100 (PBST) and blocked with 5% donkey serum (in 0.1% PBST) for 1 hours at room temperature. Primary anti-RFP (VWR, #RL600-401-379), anti-SFPQ (Abcam, #ab38148), anti-SRSF10 (Bioss, #BS-13229R), and/or anti-HuR (Proteintech, #11910-1-AP) antibody were added to slides and incubated at 4 °C overnight. Slides were washed 3 x 3 min with 0.1% PBST and then incubated with secondary antibody (Invitrogen: AlexaFluor H+L) for 1 hour at room temperature, followed by additional wash steps. Coverslips were mounted with ProLong Gold Antifade Mountant complete with DAPI (ThermoFisher).

### Confocal Microscopy and Image Analysis

All mouse histological methods were imaged on a Nikon Eclipse Ni C2 Si laser scanning upright confocal with 4 laser lines. Slides were imaged at 20x, 40x, and 60x magnification, acquired using NIS Elements software, and saved in nd2 format for further processing and analysis. Images were opened in ImageJ and subjected to basic preprocessing steps to enhance both Nav1.8+ axon and particle visibility; this included adjusting brightness and contrast and removing noise from the background by despeckling. A manual threshold was first applied to the Nav1.8+ axon channel, and a mask was generated from the selection and added as a region of interest (ROI). A manual threshold was then applied to the particle channel (displaying PLA, RNAscope, or IHC puncta), and the intensity values were adjusted to ensure clear delineation between the particles and the background. Once the appropriate threshold was set, the area of the image contained within the previously set ROI was converted into a binary mask. The Analyze Particles function was used to detect and quantify individual particles in the binary image. The analysis was performed with a size range of 1 - 100 pixels and circularity range of 0 −1.0. The results were output to a results table, which included measurements such as particle count, mean area, and standard deviation. Following automatic particle detection, the results were visually inspected to ensure accurate particle identification. If necessary, additional particles were manually added or removed from the analysis based on visual verification.

### RNA Extraction and qRT-PCR

RNA was extracted from mouse testes, liver, dorsal root ganglia, sciatic nerves, and dorsal roots by basic phenol:chloroform extraction followed by isopropanol RNA precipitation. RNA was converted to cDNA with the High Capacity RNA-to-cDNA Kit (Applied Biosystems). Template cDNA and Catsper2 primers were incorporated into a SYBR Green master mixture (Applied Biosystems), and mRNA expression was quantified using the EP Gradient s Realplex Mastercycler (Eppendorf). GAPDH was used to normalize gene expression. Each sample was assessed in duplicate and included a template-free control. All primers used were synthesized by Eurofins.

### RBP Bioinformatic Analysis

Bioinformatic RBP motif analysis of 3’UTRs were performed using the oRNAment and RBPmap databases ^39,40^. The locally translated mRNAs associated with significant functional annotations were used as the input transcript reference list. Transcripts were limited to 3’UTRs. Predicted motif score cutoff was set to 90%. RBPs found to have binding motifs with the key 3’UTRs were further analyzed for their ability to form ribonucleoprotein granules using the RPS.renlab database ^41^. Functional Annotation Analysis for RBP protein-protein interactions was performed using the STRING database ^42^. The minimum required interaction score was set to High Confidence (0.700).

### RNA Immunoprecipitation (RIP)

RNA Immunoprecipitation was performed according to the EZ-Magna RIP RNA-Binding Protein Immunoprecipitation (Millipore Sigma) protocol. In brief, on Day 5 of primary DRG neuron cell culture, cells were washed with ice-cold PBS and lysed at −80 °C overnight in Complete Lysis Buffer. Lysates were spun down to pellet cellular debris and the supernatant added to Protein A/G Beads conjugated with 5μg of primary anti-SFPQ (Abcam, #ab38148), anti-SRSF10 (MBL, #50304712), or anti-rabbit IgG (EZ-Magna Kit). Lysate-bead mixtures were incubated at 4 °C overnight with end-to-end rotation. Protein-RNA complexes were eluted from beads in a Proteinase K Buffer for 30 mins at 55 °C. RNA was further purified using phenol:chloroform:isoamyl alcohol procedure followed by absolute ethanol and salt RNA precipitation.

### RIP-Sequencing

Sequencing was performed by the DFCI MBCF:Genomics Core Facility using NovaSeq X: Low Input mRNAseq (>1ng RNA). Libraries were prepared with Takara SmartSeq v4 full length cDNA synthesis using oligo dT priming & 5’ template switching enzyme. Sequencing was performed with 40M 150bp read pairs from Illumina NovaSeq X Plus. Sequences were aligned to rat reference genome with STAR_2.7.2b^27^. RNA-seq analysis of raw counts was performed in R for sample-level quality control (principal component analysis), Differential Gene Expression testing for pairwise, group comparisons (DESeq2) ^31^, and Functional Annotation (enrichGO) ^89,90^.

### Western blot

RBP immunoprecipitations for RIP experiments were confirmed by Western Blot. Post-IP (EZ-Magna kit protocol), lysates were separated by NuPAGE 4–12% Bis-Tris SDS-Page Protein Gels and probed with anti-SFPQ (1:1000; Abcam, #ab38148), and anti-SRSF10 (1:1,000; MBL, #50304712). Bands were visualized with Donkey anti-rabbit IgG (H+L)-HRP conjugated secondary antibodies (1:10,000; Bio-Rad; 1721019) and SuperSignal chemiluminescent substrates signal (Thermo Fisher Scientific). Blots were imaged using the AI600 Chemiluminescent Imager (GE Healthcare).

## QUANTIFICATION AND STATISTICAL ANALYSIS

### Statistical Methods

All statistical analyses were performed using R v4.3.2 and GraphPad Prism v10. Differential gene expression analysis was conducted using the RStudio DESeq2 package, which applies a Wald test with FDR adjusted p-values. All histological data were quantified in Prism and are represented as mean ± standard error of the mean (SEM). Grouped data with multiple comparisons were analyzed for statistical significance using two-way ANOVA with Tukey’s multiple comparisons test. Contingency data were analyzed for statistical significance using Chi-Square. A P-value ≤ 0.05 was considered statistically significant. Data distribution was assumed to be normal, but this was not formally tested. Effect size was determined by Cohen’s *d* measurements, or the standardized difference between the means of two individual groups ((mean 1 – mean 2) / standard deviation). Statistical details of experiments can be found in the figure legends.

## ADDITIONAL RESOURCES

All raw and processed TRAP-seq and RIP-seq data from this study were deposited to Gene Expression Omnibus at GEO:GSE289550 and GEO:GSE289551 and will be made publicly available as of the date of publication.

The data are also available for browsing and analysis on the Pain-seq multi-omic data resource at http://painseq.shinyapps.io/CompartmentTRAP/, built using R Shiny (http://github.com/Renthal-Lab/CompartmentTRAPShiny).

## SUPPLEMENTAL INFORMATION

**Figure S1: TRAP-Seq Experimental Validation.** A) Endogenous L10a-EGFP expression is visible in the soma of DRG neurons and can be efficiently targeted and labeled by anti-GFP antibodies (scale bars: 40 μm (top row), 20 μm (bottom row). B). L10a-EGP protein can be immunoprecipitated from tissue lysates with anti-GFP antibodies.

**Figure S2. Functional comparison of DRG soma and axonal translatomes.** A) Schematic depicting the translatomic gene sets compared in Figure S2 (i.e. DRG soma vs. axons). B,C) Heatmaps comparing gene expression profiles within DRG soma and central axon samples (B) or peripheral axon samples (C) (n=4 samples/axon group; n=8 samples/DRG soma group). The rows of genes cluster within groups that show similar expression profiles between soma and axons, and well as groups that have higher expression in either the soma or axon samples. D,E) Gene ontology (GO) enrichment analysis visualized as a dot plot, where each dot represents a significantly enriched biological process (BP), cellular component (CC), or molecular function (MF). These dot plots show the GO terms enriched among the genes that are shared by both the DRG soma translatome and the central axon translatome (D) or peripheral axon translatome (E) (p.adj ≥ 0.05; abs[F.C] ≤ 1.5). F, G) These dot plots show GO terms that are enriched among the genes with higher expression in the central axon translatome (F) or peripheral axon translatome (G) as compared to the DRG soma translatome (p.adj ≤ 0.05; F.C ≤ −1.5).

**Figure S3. Identification of higher confidence translatomes through background filtering.** A) Schematic depicting the gene sets compared in Figure S3 (i.e. GFP+ tissue samples vs. GFP-tissue samples). B) Venn diagram detailing the number of mRNA transcripts in DRG soma, peripheral axon, and central axon translatomic datasets after background filtering (i.e. removing transcripts with expression in GFP-negative control samples). C-E). Volcano plots demonstrating the number mRNA transcripts with significantly higher expression in GFP+ TRAP samples from the DRG soma (C), peripheral axon (D), and central axon (E) as compared to GFP-negative control samples from the same tissue compartment (p.adj ≤ 0.05; fold-change ≥ 1). F) A gene concept network demonstrates the color-coded linkages between genes from the central axon translatome and significantly enriched GO terms. The size of the dot in the network correlates to the number of mRNAs in the central axon translatome associated with that GO category. The higher-confidence central axon translatome encodes for genes related to sensory perception, pain perception, and serotonin signaling. G) A gene concept network demonstrates the color-coded linkages between genes from the peripheral axon translatome and significantly enriched GO terms. The local peripheral axon translatome encodes for genes related to synaptic specialization, cilia, and cytosolic ribosomal proteins.

**Figure S4. Puromycin incorporation into DRG soma and axons.** A) Representative fluorescent immunohistochemistry (IHC) images of Puromycin and TdTomato-labeled Nav1.8+ (Nav1.8-tdT) axons in DRG soma. Scale bar = 100 μm. B) Representative fluorescent images of Puromycin and Nav1.8-tdT+ axons in sciatic nerves. Yellow, orange, and green boxes denote areas with both Puromycin + TdTomato expression, only Puromycin expression, and only TdTomato expression, respectively. Scale bar = 40 μm. B’) High magnification images of yellow, orange, and green boxes shown in (B). Scale bar = 10 μm.

**Figure S5. Validation of mRNA expression of additional transcripts from translatomic datasets.** A) qRT-PCR data demonstrating relative *Catsper2* mRNA expression in mouse testes (positive control), liver (negative control), DRG (soma), sciatic nerves (peripheral axons), and dorsal roots (central axons). All samples are normalized to GAPDH expression. Data are represented as mean ± SEM. B) Representative 20x fluorescent images of *Catsper2* mRNA puncta (visualized by RNAScope *in situ* hybridization) and TdTomato-labeled Nav1.8+ (Nav1.8-tdT) neurons in DRGs, dorsal roots, and sciatic nerves, representing DRG soma, central axons, and peripheral axons, respectively. Scale bar = 100 μm (soma) and 40 μm (axons). B’) Higher magnification images demonstrating the location and abundance of *Catspser2* mRNA puncta within Nav1.8-tdT+ soma in the ganglia, and axons in the dorsal root and sciatic nerve. Scale bar = 20 μm (soma) and 10 μm (axons).

**Figure S6. RNA Binding Protein motif analyses and localization.** A) Heatmap displaying the number of RBP motifs within the 3’UTR of mRNAs with axon-preferential translation. RBPs identified via oRNAment, RNAmap, and RPS.renlab. B-D) STRING motif analysis of the cluster of RBPs found to have 3’UTR binding sites in: both central and peripherally targeted transcripts, i.e. “shared cluster” (B), centrally targeted transcripts, i.e. “central cluster” (C), and peripherally targeted transcripts, i.e. “peripheral cluster” (D). Data are represented as protein interaction networks with a high-confidence score of 0.700. Nodes represent individual proteins and are color-coordinated by enriched GO cellular components (FDR ≤ 0.05). Edges represent protein-protein associations based on known interactions, predicted interactions, and protein homology.

**Figure S7. SFPQ and SRSF10 RIP-Sequencing.** A, B) Western blot images showing SFPQ (A) and SRSF10 (B) protein. C) Principal component analysis demonstrating the variance between SFPQ IP, SRSF10 IP, IgG IP, and paired input samples (n=4 sets/condition). C’) Principal component analysis demonstrating the variance between SFPQ IP (n=4 sets), SRSF10 IP (n=3 sets), and IgG IP (n=2 sets) samples with non-zero values only. D) Volcano plots from RIP-Seq experiments demonstrating the mRNA transcripts with significantly higher expression in IgG IP samples as compared to the input fraction from the same DRG neuron cell culture experiments. The most significant differentially expressed transcripts are labeled on the graph. E, F) Gene ontology (GO) enrichment analysis visualized as a dot plot, where each dot represents the most significantly enriched biological processes (BP), cellular compartments (CC), or molecular functions (MF) from the SFPQ IP (E) and SRSF10 IP (F) datasets (p.adj ≤ 0.05). F’) Dot plot demonstrating significantly enriched, neuron-related GO cellular compartments from the SRSF10 IP dataset (p.adj ≤ 0.05).

## Supplemental Spreadsheets

Supplemental File1 (TRAP): Excel file containing Tables S1-S12, which are too large to fit in a PDF and related to TRAP-seq experiments.

<u>Related to</u> Fig. 1:

S1. Normalized counts DRG soma samples

S2. Normalized counts for Peripheral Axon samples

S3. Normalized counts for Central Axon samples <u>Related to</u> Figs. 2<u>, S2:</u>

S4. Differential Gene Expression Analysis between Central and Peripheral Axons (CA vs. PA)

S5. Enriched GO terms: Shared Between Central and Peripheral Axons (CA vs. PA)

S6. Differential Gene Expression Analysis between Central Axons and DRG Soma (CA vs. DRG)

S7. Differential Gene Expression Analysis between Peripheral Axons and DRG soma (PA vs. DRG)

S8. Enriched GO terms: Central Axons (CA vs. PA)

S9. Enriched GO terms: Peripheral Axons (CA vs. PA) <u>Related to</u> Fig. 3

S10. Differential Gene Expression Analysis between Pain-seq and TRAP-seq <u>Related to Fig. S3:</u>

S11. Background Filtered Gene Lists (Higher Confidence) <u>Related to</u> Fig. 4:

S12. Validated Locally Translated mRNAs in Central and Peripheral Axons

Supplemental File2 (RIP):: Excel file containing Tables S13-17, which are too large to fit in a PDF and related to RIP-seq experiments.

<u>Related to</u> Fig. 6:

S13. Normalized Counts for all RIP Samples

S14. Differentially Expressed Genes in SFPQ-IP samples

S15. Differentially Expressed Genes in SRSF-IP samples

S16. Enriched GO terms: SFPQ IP

S17. Enriched GO terms: SRSF IP

---

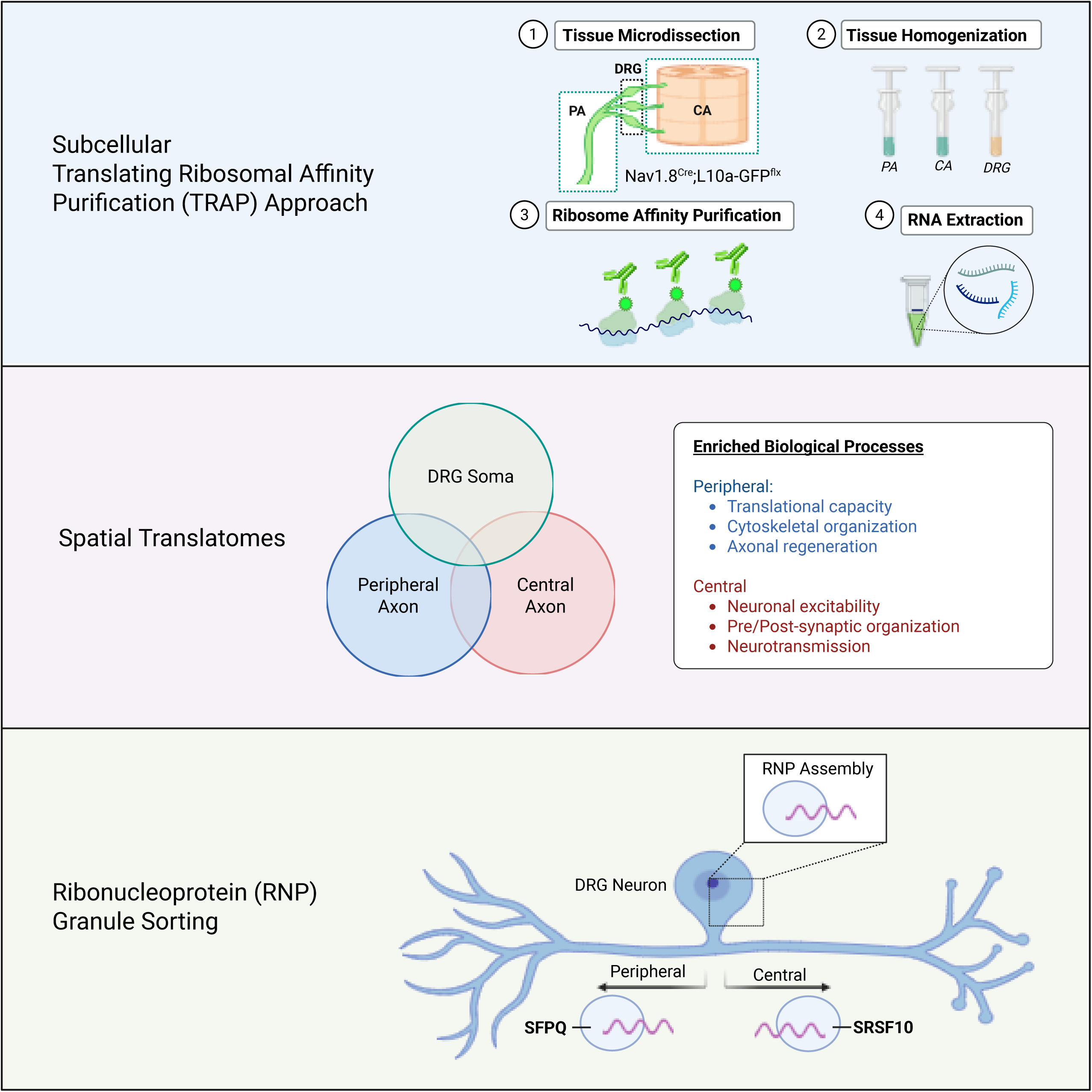

