## Supplementary figures and images for "RBP-Driven RNA Sorting and Local Translation Establish the Molecular Identities of Individual Sensory Axons"

### Supplemental Figure 1

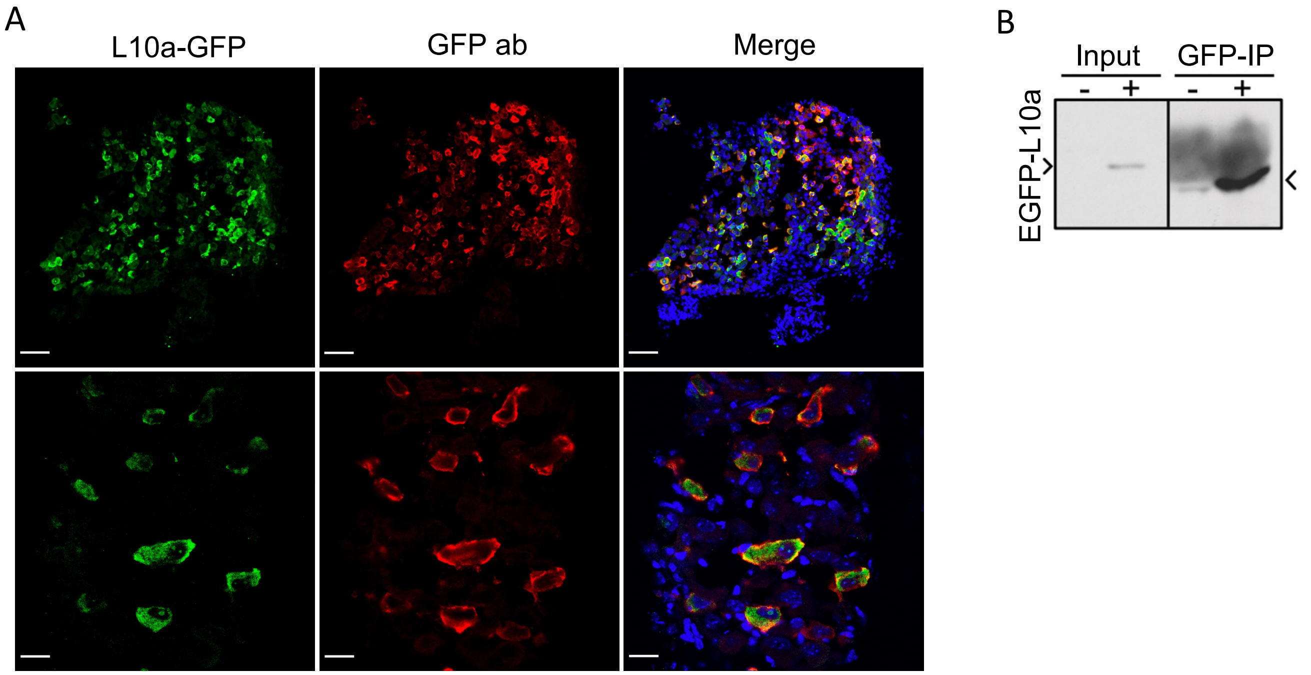


Figure S1

### Supplemental Figure 2

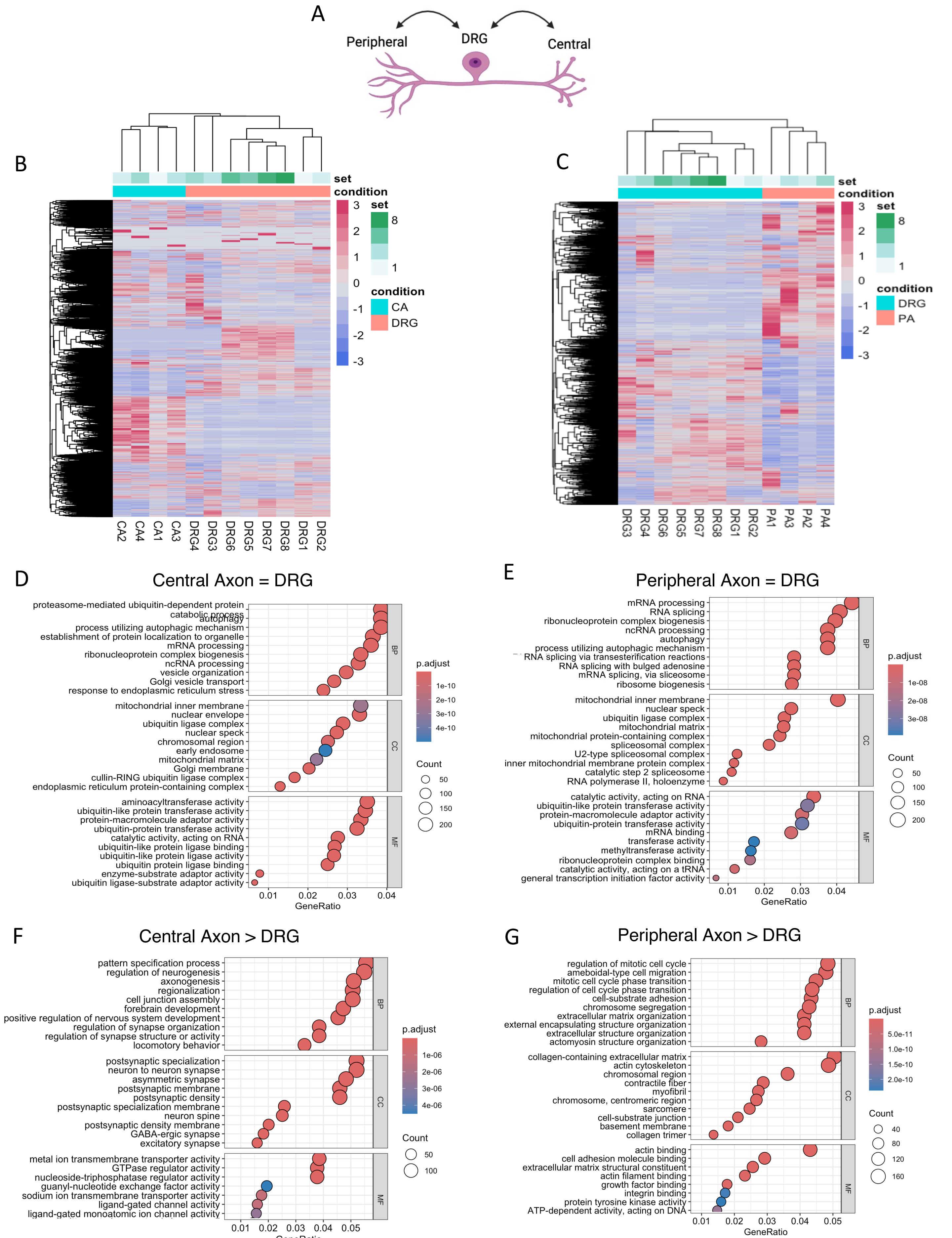

### Supplemental Figure 3

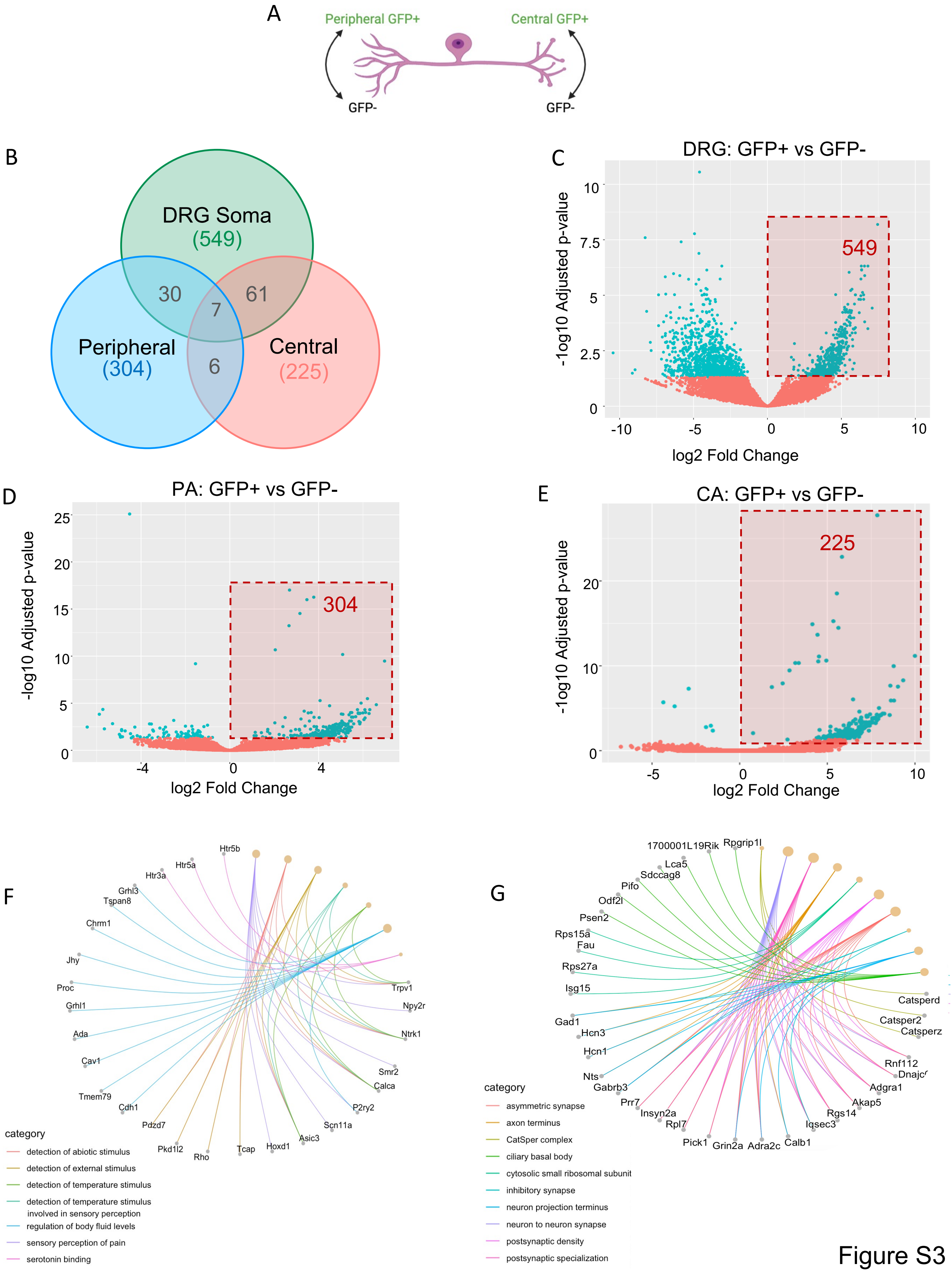

### Supplemental Figure 4

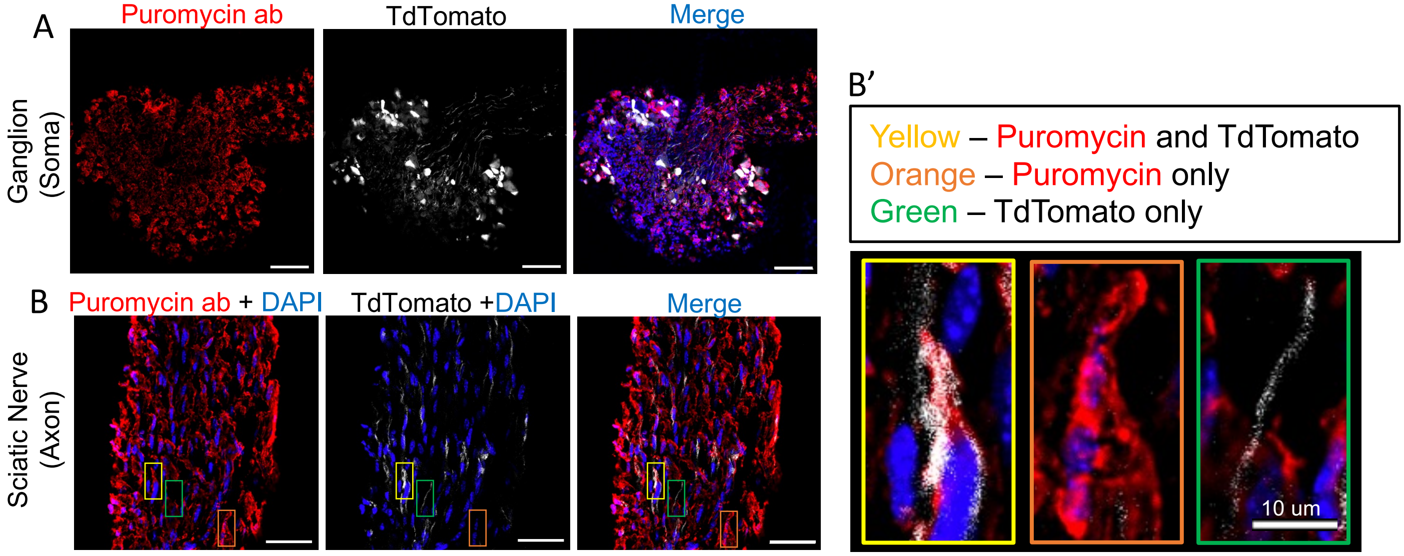

### Supplemental Figure 5

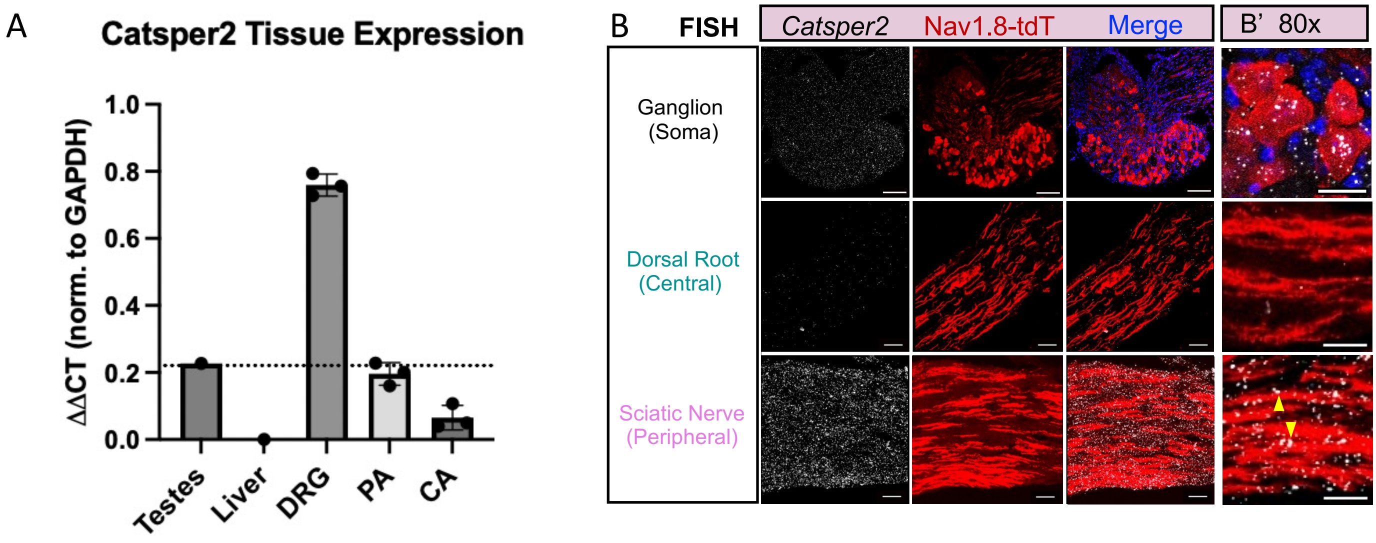

### Supplemental Figure 6

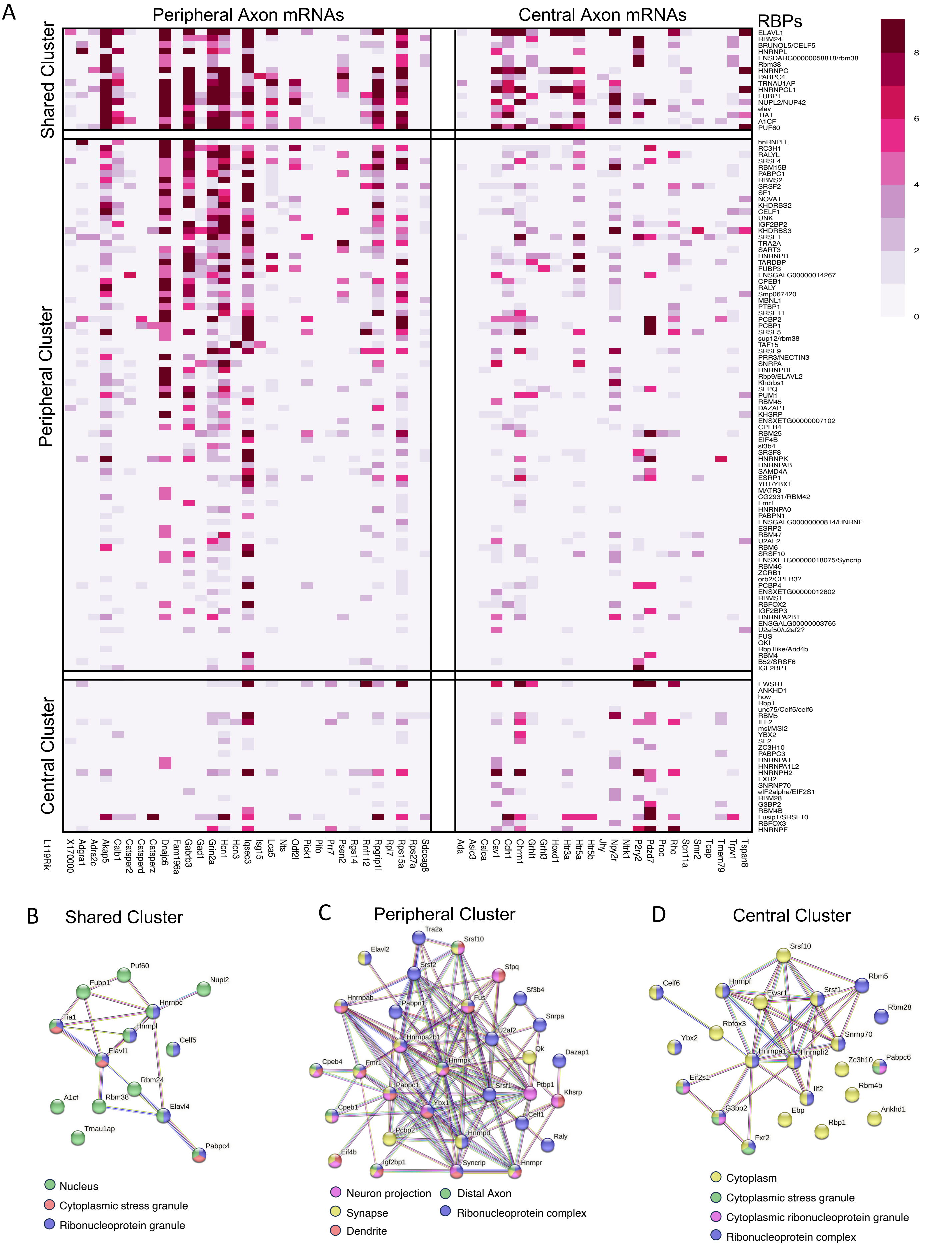

### Supplemental Figure 7

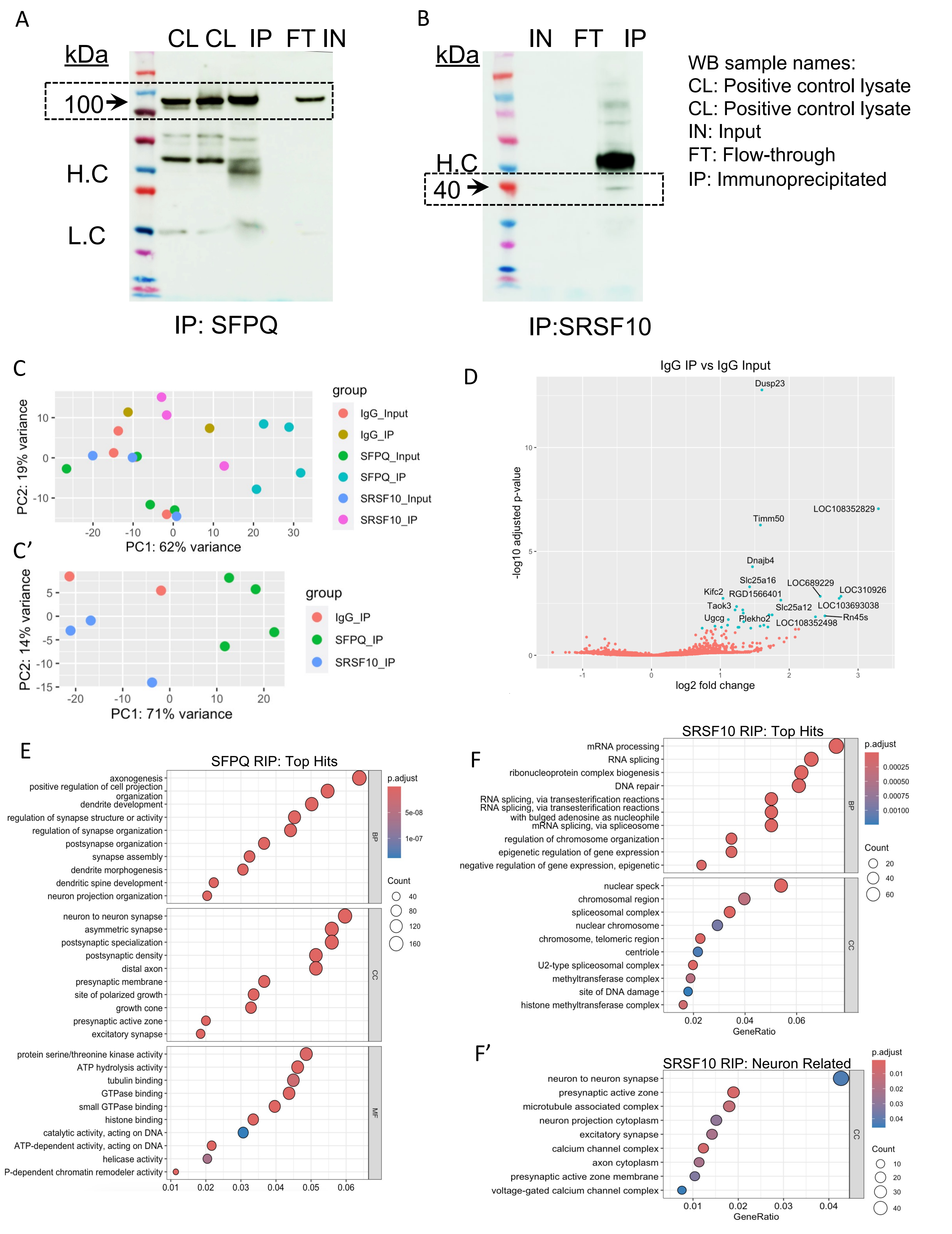
